# Utilising citizen science data to rapidly assess potential wild bridging hosts and reservoirs of infection: avian influenza outbreaks in Great Britain

**DOI:** 10.1101/2024.03.28.587127

**Authors:** Stephen H. Vickers, Jayna Raghwani, Ashley C Banyard, Ian H Brown, Guillaume Fournie, Sarah C. Hill

## Abstract

High pathogenicity avian influenza virus (HPAIV) is a rapidly evolving orthomyxovirus causing significant economic and environmental harm. Wild birds are a key reservoir of infection and an important source of viral incursions into poultry populations. However, we lack thorough understanding of which wild species drive incursions and whether this changes over time. We explored associations between abundances of 152 avian species and cases of HPAI in poultry premises across Great Britain between October-2021 and January-2023. Spatial generalised additive models were used, with species abundance distributions sourced from eBird modelled predictions. Associations were investigated at the species-specific level and across aggregations of species. During autumn/winter, associations were generally strongest with waterbirds such as ducks and geese; however, we also found significant associations in other groups such as non-native gamebirds, and rapid change in species-specific associations over time. Our results demonstrate the value of citizen science in rapid exploration of wild reservoirs of infection as facilitators of disease incursion into domestic hosts, especially in regions where surveillance programmes in wild birds are absent. This can be a critical step towards improving species-specific biosecurity measures and targeted surveillance; particularly for HPAIV, which has undergone sudden shifts in host-range and continues to rapidly evolve.

## 1. Introduction

The emergence and rapid evolution of transboundary animal pathogens poses significant economic and zoonotic risks (Clemmons et al 2021). Many transboundary animal diseases are caused by multi-host pathogens with assemblages of reservoirs of infection and bridging species that may change over time and/or vary greatly in breadth (Portier et al. 2019). Which host species are most affected can vary seasonally alongside changes in species ecology or shift rapidly as a result of pathogen evolution or diffusion into new areas (Bonneaud & Longdon 2020). Therefore, continuous monitoring of possible reservoirs of infection and bridging species is essential for designing effective control measures and developing predictive disease transmission models (Viana et al. 2014). However, widespread surveillance in wildlife is often too challenging and expensive to implement, especially for hosts with broad species distributions.

Avian influenza virus (AIV) has among the widest geographic and host range of all transboundary animal diseases (Lupiana & Reddy 2009). In 2020, the emergence of high pathogenicity AIV (HPAIV) subtype H5Nx clade 2.3.4.4b led to an epizootic of HPAIV that profoundly impacted both the poultry industry and wild bird populations, causing significant economic and environmental harm (Lewis et al. 2021, EFSA et al. 2023A). During annual HPAI epizootics in Great Britain since 2020, farm contact tracing and genetic analysis of viruses from infected poultry has strongly implicated wild birds as the likely primary source of repeated independent incursions of HPAIV infection (Byrne et al. 2023), even if the exact pathway of virus movement from wild birds to poultry cannot be proven.

Our understanding of which wild bird species are responsible for such incursions and whether this varies between epizootics due to virus evolution and heightened premise biosecurity remains relatively coarse (Hill et al. 2019, Blagodatski et al. 2021). The main wild host reservoir of infection for AIV is generally considered to be ducks and other waterfowl (Order Anseriformes, family *Anatidae*), with infection in these species resulting in disease outcomes ranging from subclinical to severe (Keawcharoen et al. 2008). In contrast, HPAIV H5Nx infection of poultry invariably results in severe disease with high morbidity and mortality rates (Ramey et al. 2022). Interestingly, recent outbreaks of H5Nx clade 2.3.4.4b have been characterised by high levels of mortality across a broader range of wild birds, including anatids, than seen previously. Infection has also shifted, likely through emergence of novel H5N1 genotypes, to cause extensive outbreaks. High susceptibility and increased virus shedding in a range of wild bird species has increased the levels of infection pressure through elevated levels of environmental contamination. This in turn has resulted in previously rarely affected groups of birds such as seabirds becoming exposed and infected (Banyard et al. 2022, Falchieri et al. 2022, Pohlmann et al. 2023). Although the host range for HPAI H5Nx are known to be broad based on challenge studies and wild-bird surveillance (Lee et al. 2017, Empress-i 2023), virulence and transmissibility appear to be highly variable across taxa. Mass mortality events attributed to H5 clade 2.3.4.4b have been recorded in several groups such as wildfowl, shorebirds, seabirds, and raptors (Abolnik et al. 2023, Adlhoch & Baldinelli 2023, Lane et al. 2023, Pohlmann et al. 2023), whereas in other groups, such as passerines, such reports are lacking. However, passerines have demonstrated moderate to high seroprevalence against older clades of HPAI H5Nx, likely indicative of past infection (Kou et al. 2005, Kaplan & Webby 2013), and detecting increased mortality in these less conspicuous species is likely to be more difficult. Importantly, commonly tested species that contribute to the long-distance spread of HPAIV internationally (particularly wildfowl (Order Anseriformes) and gulls (Family *Laridae*); Kim et al. 2009, Hill et al. 2022) may not necessarily be sufficient to explain incursions onto premises. Instead, there may be additional species acting as local amplifiers of viruses, which could be critical in facilitating viral incursion into premises.

Understanding species-specific patterns of spillover from wild to domestic birds is difficult due to obstacles in effective monitoring of HPAIV in potential wild bird hosts, and the changes in species assemblages and abundance across the annual cycle. Despite significant advances in testing protocols (James et al. 2022, Slomka et al. 2023), surveillance of HPAI outbreaks in wild birds can be severely limited by financial constraints, detection and sampling of carcasses, and requirements for high biosecurity testing facilities (Hill et al. 2018). Testing may therefore be heavily skewed towards species with overt clinical symptoms or for which carcasses are more easily detectable. For example, while all suspected kept bird premise cases of HPAI in Great Britain are tested, only a relatively small subset of reported wild bird carcasses suspected of being HPAI cases can be tested (DEFRA 2023C). Therefore, methods to refine targeted surveillance of HPAIV are essential to maximise disease control efforts.

One way to potentially achieve better targeted surveillance is by assessing associations between the abundance of potential reservoirs of infection or bridging species and confirmed HPAIV infection in premises housing domestic poultry or other captive birds (henceforth referred to collectively as “captive avian premises”), on the assumption that there will often be a higher incidence of cases where one or more reservoirs of infection or bridging species are at higher abundance. While this approach does not rely upon *a priori* knowledge of a single host species, as several species can be assessed concurrently, it does rely on an accurate understanding of spatiotemporal patterns of possible host species abundance across the region of interest. Modelling spatiotemporal patterns of wild host abundance at national scales is complex, because it requires a large amount of observational data and computationally intensive analyses (Fink et al. 2013). However, citizen science initiatives alongside advances in computational power and novel analytical approaches that account for biases in semi-structured data collection procedures can enable rapid advances in our understanding of species abundance and distributions. The citizen science initiative eBird launched by the Cornell Lab of Ornithology (CLO) and the National Audubon Society in 2002 has amassed over 1 billion bird observations globally and currently produces estimates of the full annual cycle distributions, abundances, and environmental associations for over 2000 species (Fink et al. 2022). With these advances, it is now possible to rapidly assess the associations between the spatial distribution of wild bird populations and HPAI in poultry at larger scales than previously possible.

As many different wild bird species may be involved in HPAIV transmission, with varying levels of interaction with each other and captive avian premises, assessing species-level drivers of virus incursions into captive flocks can be challenging. Using eBird-modelled species abundance estimates that are publicly available for a broad selection of wild avian species, we investigate spatial associations between species-specific wild-bird abundance and patterns of HPAI cases in captive avian premises across Great Britain. This will enable us to identify potentially under-prioritised key species, or species groups, that may be facilitating the incursion of HPAI into captive avian premises.

The use of citizen science recording schemes such as eBird benefits from being non-reliant on any field- or laboratory-led surveillance in potential wild viral hosts, and as such can be rapidly deployed against pathogens with wild reservoirs of infection in areas with limited wild host testing capacity. Continued monitoring of associations between wild hosts and detected outbreaks in ‘captive avian’ premises could also prove useful as an early-warning sign of a possible virus host range expansion into a new wild species that is not currently subject to surveillance.

## 2. Methods

We used publicly available model-predicted relative abundance distributions (RADs) of wild birds to explore how wild bird abundance is associated with HPAI cases in premises across Great Britain between 19^th^ October 2021 and 20^th^ January 2023. RADs are built upon data from eBird, a global community science bird monitoring program administered by The Cornell Lab of Ornithology (Strimas-Mackey et al. 2022; Fink et al. 2023).

### A. Data processing

RADs were retrieved from eBird Status and Trends using R package ebirdst (Strimas-Mackey et al. 2022). These RADs provided estimates for the full annual cycle at weekly intervals, modelled on the year 2021, across a regular grid which we use at a resolution of 26.7km^2^. Predicted RADs are derived from an ensemble modelling strategy based on the Adaptive Spatio-Temporal Exploratory Model (AdaSTEM; Fink et al. 2013). For full RAD methodology see Fink et al. (2020). RADs that include Great Britain are currently available for 256 species, each with seasonal quality ratings assessed between 0 (lowest quality) and 3 (highest quality). We consider only those species with seasonal quality rating scores of 2 or 3 during the modelled period (to exclude predictions with low confidence), and a sum of relative weighted abundance scores within the relevant time period of >1 (to exclude uncommon and rare species). This produced a final pool of 152 species. Seasonal quality rating scores are assigned based upon expert human review, whereby a rating of 0 implies the modelled predictions failed review and ratings of 1–3 correspond to increasing levels of extrapolation and/or omission.

During our study period, there were 312 HPAI cases across commercial (*n* = 216), backyard (*n* = 70), mixed (*n* = 12), and miscellaneous (e.g., rescue centres, zoos, etc.; *n* = 14) premises in Great Britain (referred to here collectively as premises; Fig 1). Case data on infected premises housing captive birds was obtained from the Animal and Plant Health Agency (APHA), who lead diagnostic surveillance for AIV in Great Britain (Byrne et al. 2023). Through visual inspection of epidemic curves, cases were split into three distinct EP’s. EP 1 covered 19^th^ October 2021 – 9^th^ February 2022 coinciding with a large peak in cases, EP 2 covered 10^th^ February 2022 – 16^th^ August 2022 during which cases occurred infrequently, and EP 3 covered a large peak in cases between 17^th^ August 2022 – 20^th^ January 2023 (Fig 1A).

**Fig 1.**
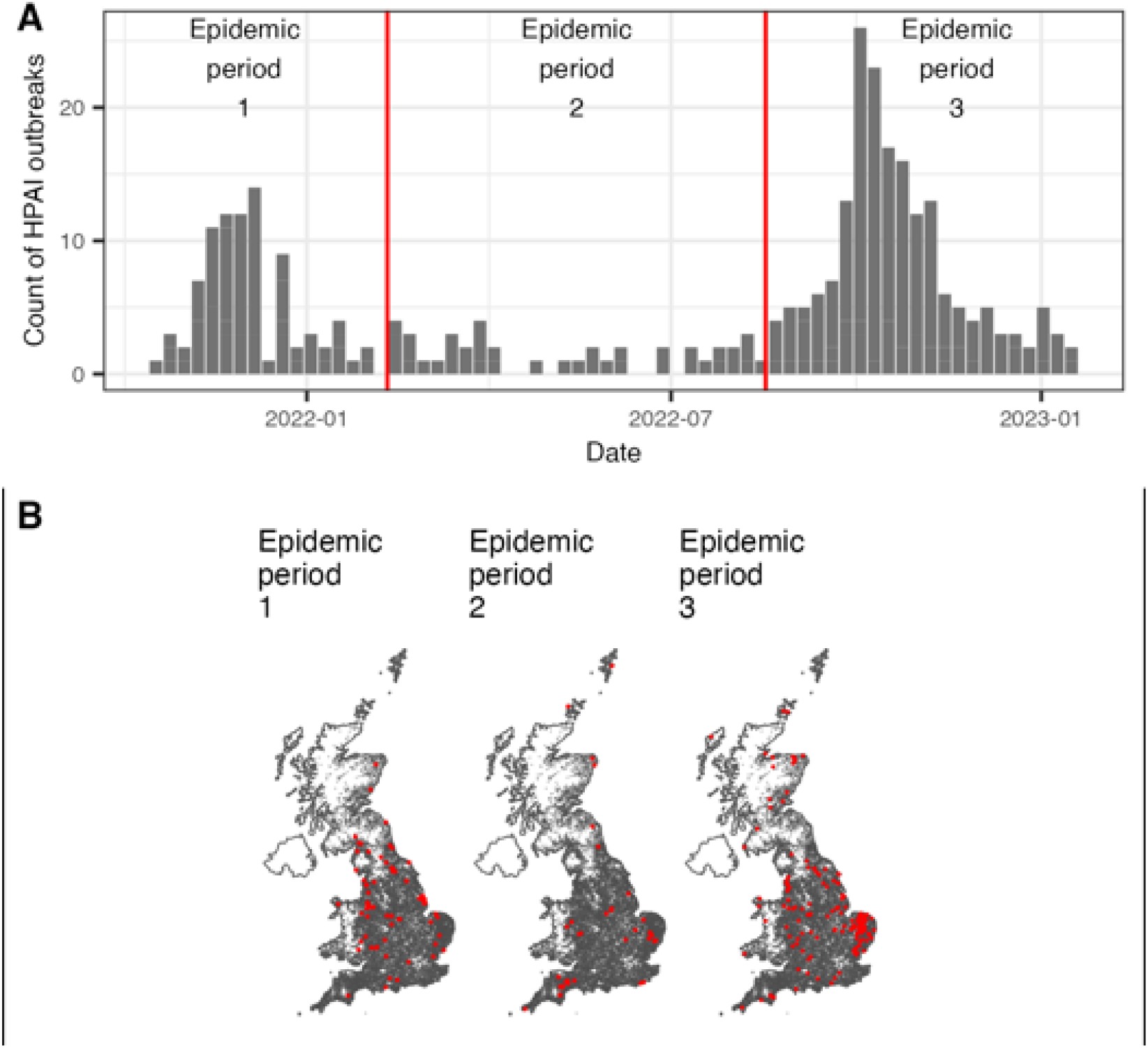
HPAI cases in captive avian premises in Great Britain between 19^th^ October 2021–20^th^ January 2023. **A)** Epidemic curve split into three distinct epidemic periods (EP1 = 19^th^ October 2021 – 9^th^ February 2022; EP2 = 10^th^ February 2022 – 16^th^ August 2022; EP3 = 17^th^ August 2022 – 20^th^ January 2023). Number of cases per period were 86, 40, and 186, respectively. **B)** Map of cases (red points) against a map of premises (black points) as listed in the Great British Poultry Register (jittered here to maintain anonymity).

To account for changes in each species’ relative abundance and HPAI case intensity across an EP, we weighted the weekly RADs by multiplying each cell value by the proportion of HPAI cases that occurred nationally within that week of the EP. We then sum these weekly weighted values to produce a single weighted relative species abundance for each grid-cell for the relevant period (S1 Fig). Weighted RADs therefore represented how the species abundance varied across the UK during the period, with higher weighting given to weeks with more premise cases.

In order to control for greater case likelihood being expected in areas with higher farm and poultry density, we used information on location and stock numbers of premises obtained from the Great Britain Poultry Register (GBPR). As reporting to the GBPR is non-compulsory for flocks of fewer than 50 birds, we only used data from those farms with 50 or more birds (19,680 farms). Farm locations and reported stock counts were rasterised across a regular grid at a resolution of 26.7km^2^ to match wild bird RAD data, giving a count of farms and sum of stock numbers at the grid-cell level (S1 Fig).

As an alternative to wild bird RAD data, we also explored associations of premise cases with PCR-confirmed wild bird HPAI cases within each period. Spatial data on individual HPAI cases in wild birds was sourced from the Empres-i open-access database (Martin et al. 2007). Case locations were rasterised across a regular grid at a resolution of 26.7km^2^ to match wild bird RAD data to give a count of cases at the grid-cell level within each period (S2 Fig).

### B. Primary data analysis

We fit spatial generalised additive models (GAMs; package *mgcv*; Wood 2011) with a binomial error family and a logit link function to test the effect of relative species abundance on premise cases at a 26.7km^2^ grid-cell level. Our dependent variable (proportion of all infected premises over the epidemic period which were located in each grid cell) was included as a two-column count matrix of infected premises and uninfected premises in order to perform weighted regression using the total number of premises within the grid-cell as weights. Explanatory variables of weighted species abundance and average stock number per premise were included as linear terms, which were scaled prior to model fitting. To account for spatial autocorrelation between neighbouring grid-cells, we also include a two-dimensional splined variable of latitude and longitude, fit with an isotropic smooth on the sphere. The default basis dimension (k) value of -1 was used, and Generalised Cross-Validation was used to estimate smoothness parameters. A separate model was fit for each species and epidemic period where sufficient species abundance was present (defined as species with a sum of weighted relative abundance across all grid cells within that period greater than 1). Models that did not converge were removed from the model pool.

In addition to this, we also fit models where the count of wild bird HPAI cases at the grid-cell level was used instead of weighted relative species abundance. This resulted in a total of 389 independent models. To account for multiple testing, p-values for species-specific models were adjusted using two methods. We first used the false discovery rate method which accounts for the expected proportion of false discoveries amongst the rejected hypotheses (Benjamini & Hochberg 1995), and second the more conservative Holm-Bonferoni method (Holm 1979).

### C. Post-hoc analysis

We fit linear models (LMs) in order to assess group-level effects across our species-specific relative abundance associations. The linear slope coefficient of relative abundance from each of the species-specific independent models was used as the dependent variable, and the model was weighted by the reciprocal of the squared standard error of this slope coefficient. Categorical explanatory variables included the Epidemic Period (1, 2, or 3), species grouping (see below), and the two-way interaction between these terms. Several models of the same structure were run with varying coarseness of the species grouping variable. Our coarsest grouping categorised species into three ecological sets of species – *Landbirds*, *Seabirds*, and *Waterbirds*, following classifications described in Geen et al. (2019). We then assessed group-level patterns based upon commonly used colloquial species groups. These groupings were chosen to reflect similarity in species behaviour, habitat-use, and phylogeny, which may all influence a species likelihood of interacting with premises and being a HPAIV carrier. Some groupings were however poorly represented as not all species present in Great Britain are modelled by eBird or because model outputs were poorly supported (i.e., low seasonal quality rating scores). We measured grouping coverage by comparing against the BOU British list (excluding vagrants; BOU 2023; S5 Table).

To assess correlation between species-specific slope coefficients for the effect of species abundance across EP’s, Pearson’s product moment correlation was used. We assessed group-level estimates of change in species-specific slope coefficients for the effect of species abundance between EP’s 1 and 3 using a general linear model, change in slope coefficient from EP1 to EP3 explained by a single explanatory variable of species grouping. A separate model was run for each level of coarseness in species groupings.

All statistical analyses were performed with R 4.3.0 (R Core Team 2023).

## 3. Results

To assess which wild bird species are associated with HPAIV incursions into captive-avian premises, we modelled spatial associations between the abundance of 152 wild bird species and cases of HPAIV in premises during three distinct epidemic periods (EP’s) between October 2021 and January 2023 (Fig 1). We use “case” to refer to any captive-avian premise in which 1 or more captive birds are confirmed to have HPAIV by PCR and/or genetic sequencing; and “premises” to refer collectively to commercial, backyard, and miscellaneous (e.g., rescue centres, zoos, etc.) premises where captive birds are kept. Most cases were, however, reported from commercial premises (*n* = 216, 69.2%). EP’s 1 and 3 (19^th^ October 2021 – 9^th^ February 2022 and 17^th^ August 2022 – 20^th^ January 2023, respectively) covered large peaks in cases (*n* = 86 and *n* = 186, respectively) with a broad spread of cases throughout much of Great Britain. Conversely, EP 2 (10^th^ February 2022 – 16^th^ August 2022) had lower case incidence (*n* = 40) and less ubiquitous spatial occurrence of cases, with regions such as northern and central-southern England having no cases despite relatively high farm density (Fig 1B, S1 Fig).

We assessed associations aggregated across groups of ecologically and phenotypically similar species (to minimise type-1 errors; S1 Table), and at the individual species level (for full species-specific model coefficients see S1 Data). Groupings of different species were chosen to reflect similarity in species behaviour, habitat-use, and phylogeny which may influence either a species likelihood of interacting with a premise and/or being a HPAIV reservoir of infection or bridging species. Species-specific abundance associations with HPAIV cases were obtained from models in which the effects of total poultry stock within the area were controlled for. Linear slope estimates for the effect of total poultry stock varied between models but were always positive (average slope estimates (95% CI): EP1 = 15.1 (14.6–15.5), EP2 = 23.1 (22.5–23.7), EP3 = 2.9 (2.8–3.1)), indicating areas with higher total poultry numbers had a greater proportion of infected premises during an epidemic period.

### A. *Species groupings*

Our most coarse grouping considered 3 species groups: *landbirds*, *seabirds*, and *waterbirds* as described in Geen et al. (2019). *Waterbirds* showed consistent group-level significant positive associations with HPAI cases across EPs 1 and 3 (Fig 2; S2 Table). This indicates that areas with high *waterbird* abundance were likely to have a higher proportion of premises experiencing HPAI cases during these epidemic periods than areas with lower abundance for this group. *Seabirds*, in contrast, had positive associations in all EPs but only showed a significant positive association within EP 3, indicating an increased importance for this group during this later period (Fig 2, S2 Table). *Landbirds* showed no significant positive association in any EP (Fig 2, S2 Table), and this was driven in part by a lack of consistency in the direction of associations between species or species groups within this wider grouping. We subsequently considered subgroups of species within our *landbird*, *seabird*, and *waterbird* groupings based upon taxonomy.

**Fig 2.**
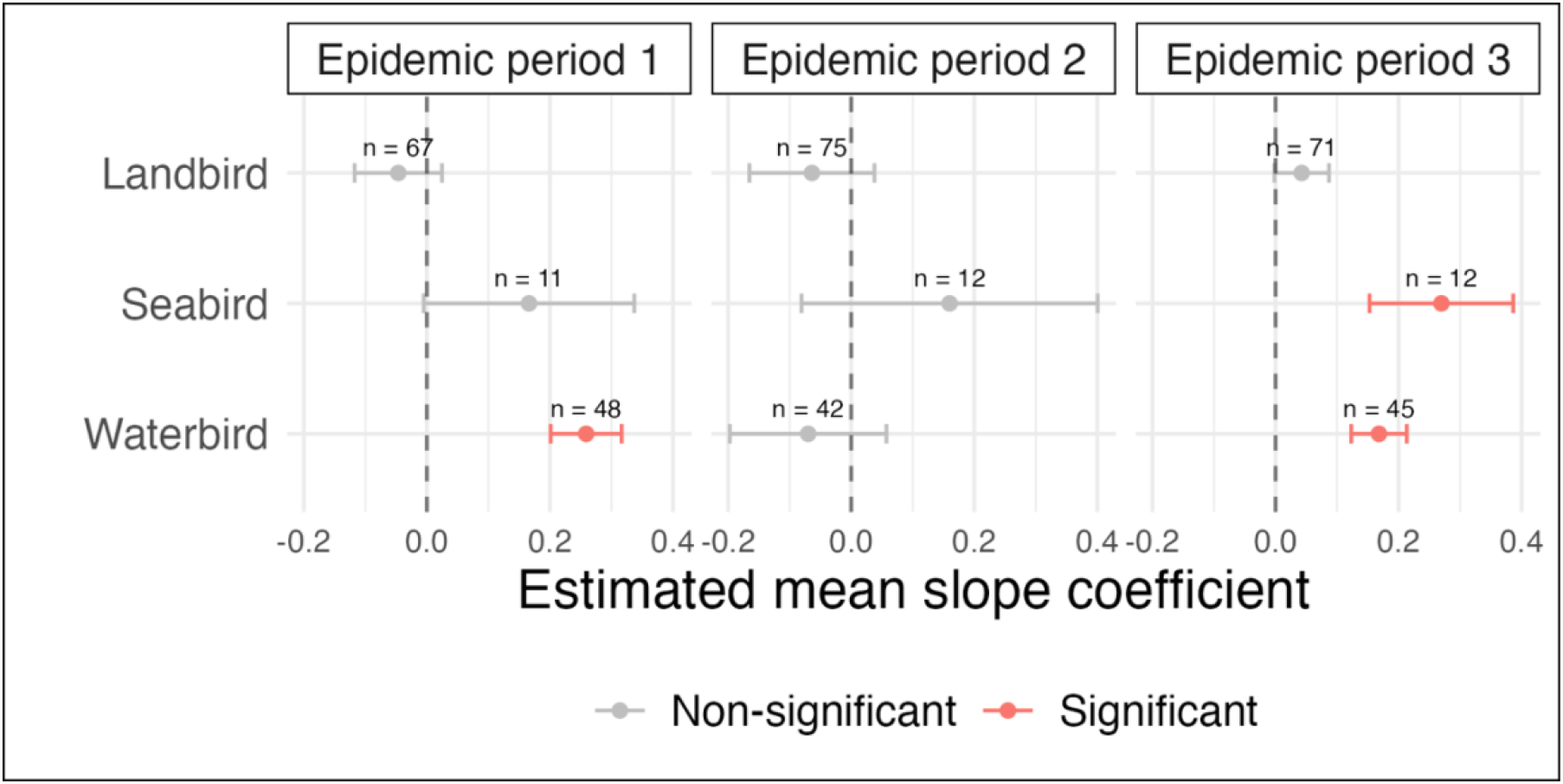
Estimated first-order group-level means for linear model coefficients describing effects of weighted relative species abundance on probability of HPAI cases in premises. Estimated means derived from a GLM assessing group effects across epidemic periods (EP1 = 19^th^ October 2021 – 9^th^ February 2022; EP2 = 10^th^ February 2022 – 16^th^ August 2022; EP3 = 17^th^ August 2022 – 20^th^ January 2023). Values above points indicate the number of species within each grouping. Error bars indicate the 95% confidence interval.

### B. *Landbirds*

Despite no significant association for *landbirds* as a wider group in any EP, significant associations were found in some subgroupings (Fig 3, S3 Fig, S3 Table, S4 Table). HPAI cases in premises during EP 1 had a significant positive association with *birds of prey* (families *Strigiformes, Accipitridae*, and *Falconidae*), although this was limited to *diurnal raptors* (families *Accipitridae* and *Falconidae*), which also had a significant positive association during EP 3. The subgroup *Passerines* (order Passeriformes) showed a significant negative association, however there was inconsistency amongst subgroups. Subgroupings such as *tits* (family *Paridae*), *flycatchers & chats* (family *Muscicapidae*), and *thrushes* (family *Turdidae*) showed significant negative associations, whereas *sparrows* (family *Passeridae*) had a significant positive association with HPAI cases in premises. This later significant positive association is of note, as some sparrows such as House Sparrow *Passer domesticus* may be regular visitors to poultry farms (Sánchez-Cano et al. 2024), may nest or roost in eaves of barns and other farm buildings, and therefore may have elevated potential to act as direct sources of infection. Furthermore, this significant positive association with *sparrows* is also observed during EP3.

**Fig 3.**
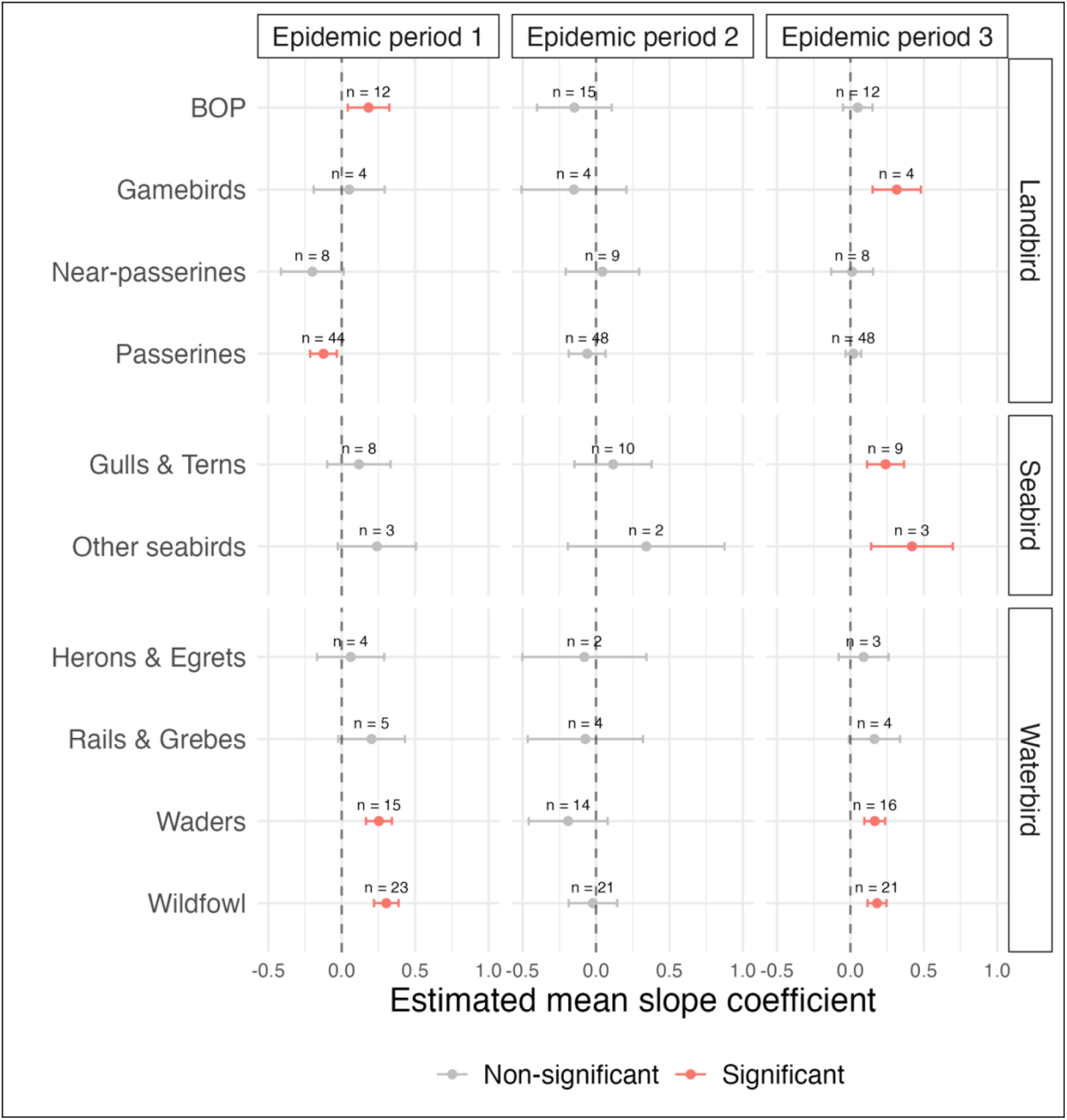
Estimated second-order group-level means for linear model coefficients describing effects of weighted relative species abundance on probability of HPAI cases in premises. Estimated means derived from a GLM assessing group effects across epidemic periods (EP1 = 19^th^ October 2021 – 9^th^ February 2022; EP2 = 10^th^ February 2022 – 16^th^ August 2022; EP3 = 17^th^ August 2022 – 20^th^ January 2023).

During EP3, *gamebirds* (family *Phasianidae*) gained a significant positive association with HPAI cases in premises (Fig 3, S3 Table). This effect was strongest amongst *non-native gamebirds*, which are bred and released in large numbers during the late-summer and early-autumn period that is included within EP 3 (Madden 2021, S3 Fig, S4 Table). In *native gamebirds*, the significant positive association was primarily driven by Grey Partridge *Perdix perdix*, which are also bred and released in some areas as part of species recovery projects, albeit in far smaller numbers (Ewald et al. 2022; S4 Fig). As was the case in EP 1, *passerine* subgroupings displayed inconsistency; *flycatchers & chats* had a significant negative association with cases, whereas other groups such as *buntings* (family *Emberizidae*) and *larks, pipits, & wagtails* (families *Alaudidae* and *Motacillidae*) had a significant positive association (S3 Fig, S4 Table).

There was no significant association for near*-passerines* (families *Columbidae*, *Picidae*, *Psittaculidae*, and *Alcedinidae*) during any EP, nor were there any significant associations amongst any of the *landbird* subgroupings during EP2 (Fig 3, S3 Fig, S3 Table, S4 Table).

### C. *Waterbirds*

*Wildfowl* (order Anseriformes) showed significant positive associations across EP 1 and EP 3 (Fig 3; S3 Table). All subgroupings had positive associations, although not all are significant (S3 Fig; S4 Table). *Dabbling ducks* (subfamily *Anatinae*) were significant in both periods, whereas *diving ducks* (subfamily *Aythyinae* and tribe Mergini) were only significant in EP 3 (S3 Fig; S4 Table). Other subgroupings were non-significant; however, there were significant positive species-specific associations in highly abundant, and therefore potentially important, species such as Greylag Goose *Anser anser* and Mute Swan *Cygnus olor* (S4 Fig). *Waders* (suborders Charadrii and Scolopaci) also showed significant positive associations in both EP 1 and 3 (Fig 3; S3 Table), driven by subgroups *plovers* and *sandpipers* (families *Charadriidae* and *Scolopacidae*) in both periods (S3 Fig; S4 Table). Across the other subgroups within *waterbirds,* namely *herons & egrets* (family *Ardeidae*) and *rails & grebes* (families *Rallidae* and *Podicipedidae*), there were no significant associations during any of the EPs (Fig 3; S3 Fig).

### D. *Seabirds*

Despite having a positive association across all EPs, *seabirds* were only found to have a significant association in EP 3. This broad grouping generally had a poorer species coverage in the available modelled eBird abundance datasets compared to *waterbirds* and *landbirds* (S5 Table). Many of the more substantial sub-groupings, such as Auks (family *Alcidae*), could not be assessed. Here, *seabirds* are predominantly represented by the *gulls & terns* (family *Laridae*) subgrouping, which had a significant positive association with premise cases in EP 3 (Fig 3; S3 Table). This significant association also remained when this subgrouping was split into *gulls* (subfamily *Larinae*) and *terns* (subfamily *Sterninae*) separately (S3 Fig; S4 Table).

The *other seabirds* grouping was also significantly positive in EP 3 (Fig 3; S3 Table). This grouping only represents a limited range of seabird species (mainly divers), and the only significant subgrouping was *divers* (family *Gaviidae*) in EP 3 (S3 Fig; S4 Table). Seabird groups such as Auks and species such as the Northern Gannet *Morus bassanus* or Great Skua *Stercorarius skua*, which have been linked to mass HPAI outbreaks in wild birds in Great Britain (Banyard et al. 2022, Lane et al. 2023), were not represented in the modelled eBird dataset.

### E. *Species-specific associations*

Generally, for species assessed in multiple EP’s, species-specific associations were poorly correlated between subsequent periods (EP1–EP2: cor = -0.075 (95% CI: -0.264–0.119), *df* = 102, p = 0.447; EP2–EP3: cor = 0.113 (95% CI: -0.085–0.303), *df* = 98, p = 0.263; Fig 4A & B), indicating a potential change in viral dynamics and potential host species during the summer period of low HPAIV incidence compared to the winter periods of high incidence. Conversely, species-specific associations were significantly correlated between EPs 1 and 3, which are both characterised by large epidemic peaks (EP1–EP3: cor = 0.691 (95% CI: 0.581–0.776), *df* = 112, p < 0.001; Fig 4C).

**Fig 4.**
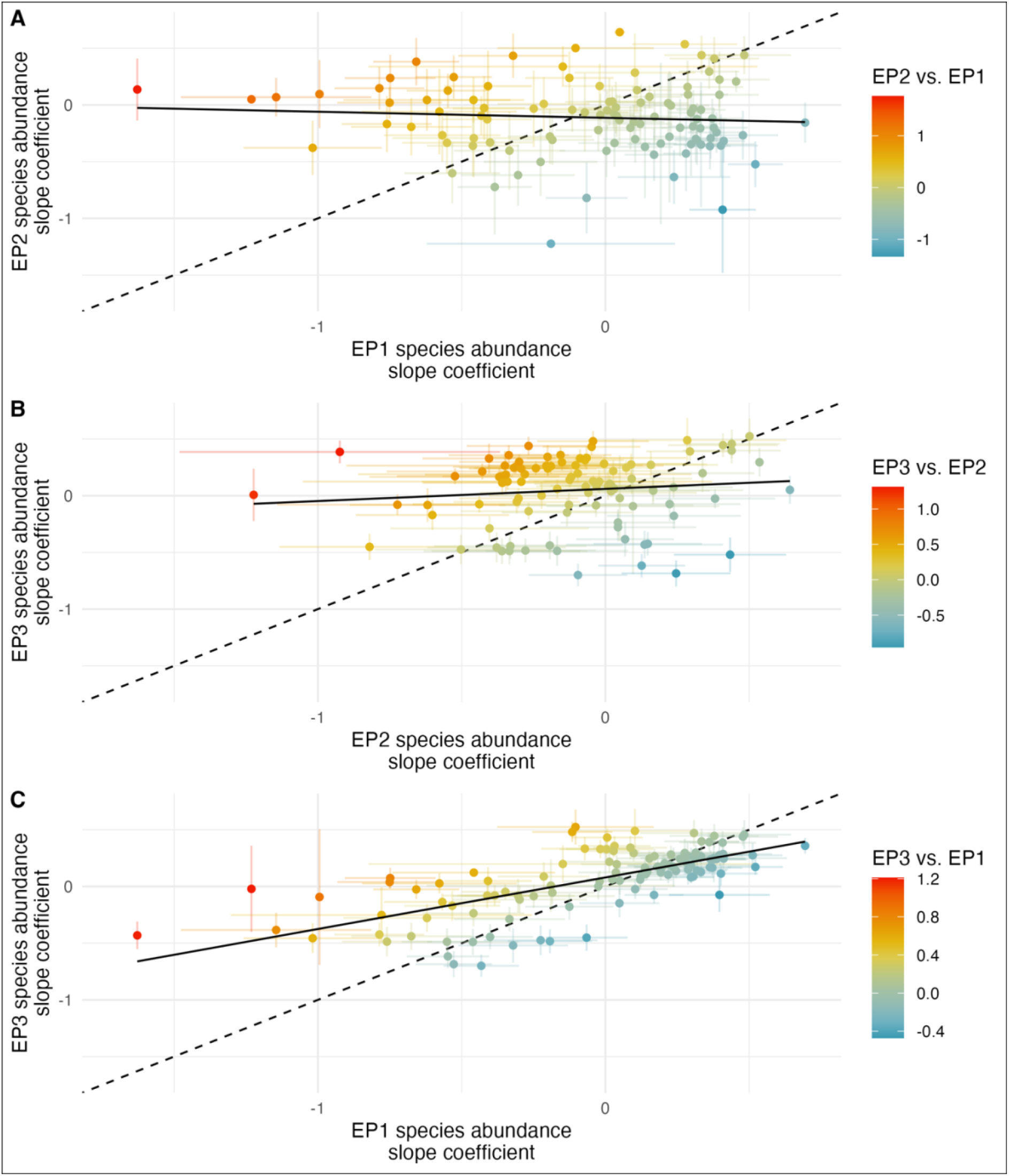
Change in species-specific slope coefficient for the effect of species abundance on case probability in premises across Great Britain. **A)** Epidemic period (EP) 2 compared to EP1. **B)** EP3 compared to EP2. **C)** EP3 compared to EP1. Dashed black line has a slope of 1 and indicates the relationship if there was no change between EPs. Solid black line is the actual line of best fit through points based upon a linear model. Error bars around points indicate standard error.

Despite significant positive correlation between species abundance slope coefficients in EP 1 and 3, there were also distinct outliers where coefficients shifted between EPs. These outliers are of potential interest, as they may represent changes in typical viral host range. Both *landbirds* and *seabirds* had significant increases in effect sizes in EP3, as did subgroups *gulls & terns* and *passerines*. All other *landbird* and *seabird* subgroupings had non-significant increases. *Waterbirds*, and its subgroupings, all showed small non-significant decreases (Fig 5, S6 Table).

**Fig 5.**
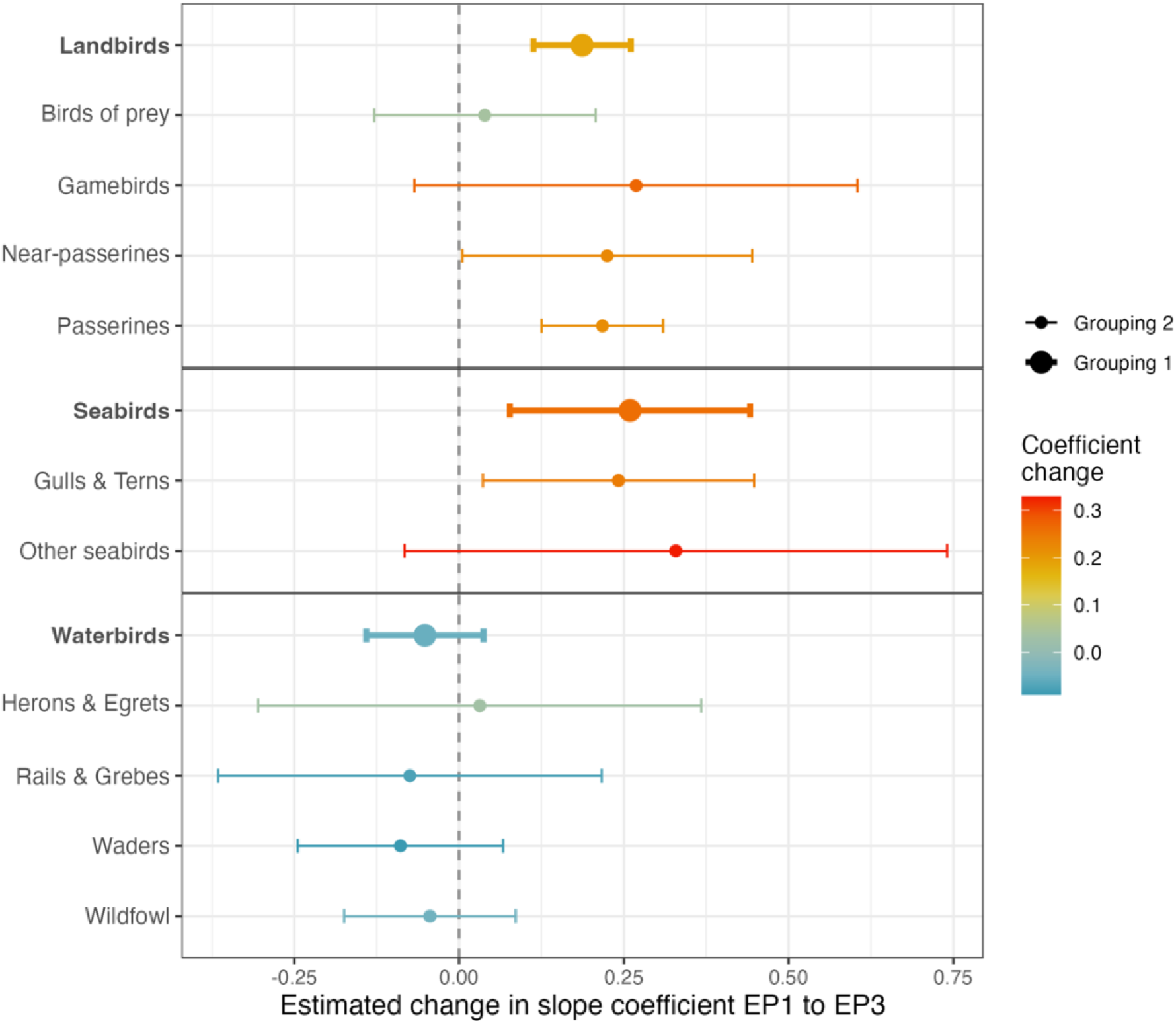
Estimated means for change between epidemic periods in the linear model coefficients describing effects of weighted relative species abundance on probability of HPAI cases in premises. Estimated means derived from a GLM assessing group effects (separate model for grouping 1 and grouping 2). Change is calculated as slope coefficient in epidemic period 3 minus slope coefficient in epidemic period 1. Bold text and thicker error bars indicate models under the coarsest species grouping (Grouping 1).

Due to the potential for type 1 errors (i.e., false positives) with multiple testing across many independent models, single-species relationships may risk overinterpretation of model results, as some significant positive spatial associations may arise by chance alone. Without correcting for multiple testing, significant positive associations were found in 41 (32.3%), 8 (6.2%), and 59 (45.7%) species across the three EPs, respectively (S4 Fig). To account for this, p-values (and therefore significance) can be adjusted. After applying the false discovery rate method (FDR) to account for the expected proportion of false discoveries amongst the rejected hypotheses (Benjamini & Hochberg 1995), significant positive associations were found in 37 (29.1%), 1 (0.8%), and 54 (41.9%) species across the three EPs, respectively (S4 Fig). However, the alternative and more conservative Holm-Bonferoni (Holm 1979) adjustment method left only 22 (17.3%), 1 (0.8%), and 27 (20.9%) species significant across the three periods, respectively (S4 Fig).

Significant negative associations were also found in 25 (19.7%), 5 (3.9%), and 21 (16.3%) species across the three EPs, respectively (S5 Fig; FDR: 23 (18.1%), 0 (0%), 20 (15.5%); Holm-Bonferoni: 10 (7.9%), 0 (0%), 16 (12.4%)). Negative associations were, however, generally of lower magnitude and were also relatively uncommon in EP1 and EP3, where we had greater statistical power due to larger numbers of HPAI cases.

### F. *Associations with wild-bird HPAI cases*

Our methods do not require data on HPAIV detections in wild birds. To explore whether premise infections can be predicted by use of wild bird abundance data alone as effectively as or more effectively as when using data on confirmed cases in wild birds, we tested whether PCR-diagnosed AIV-positive wild bird cases were associated with infected premises. The number of HPAI confirmed cases in wild birds fell across the EPs, with 425, 163, and 96 cases reported, respectively (Empres-I database; S6 Fig); however, this was likely a substantial under-representation of actual wild bird HPAI-mortality rates across these periods. Wild bird cases were positively associated with infected premises across all EPs, but the association was only significant within EPs 1 and 2 (EP 1 (winter 2021/22): Estimate = 0.050 ± 0.019SE, p = 0.009; EP 2 (spring/summer 2022): Estimate = 0.641 ± 0.112SE, p < 0.001; EP 3 (autumn/winter 2022/23): Estimate = 0.051 ± 0.123SE, p = 0.679; S4 Fig).

### G. *Model performance*

The species-specific models fitted to EPs 1 and 3 generally performed better than models fitted to EP 2, with a broadly similar median deviance explained across the species assessed in these periods (EP1 = 0.341 ± 0.002SE, *n* = 127; EP2 = 0.242 ± 0.002SE, *n* = 130; EP3 = 0.373 ± 0.002SE, *n* = 129; S7 Fig). The poorer model performance in EP 2 is likely driven by the comparatively low number of cases of infected premises (*n* = 40; S6 Fig), combined with the occurrence of a weaker spatial clustering of cases. Furthermore, this poor model performance is also likely to be a major contributing factor to the greater dissimilarity between species-specific associations during EP2 across closely related species.

## 4. Discussion

Gaining an understanding of which wild species may be key reservoirs of infection or bridging species can be extremely challenging, particularly in dynamic, multi-host systems. Our study demonstrates how publicly available citizen science outputs of wild species abundance can be a powerful tool for exploring associations between wild host abundance and disease cases in domesticated species, without relying upon targeted surveillance across potential wild host populations. Specifically, we assessed which avian abundance distributions from eBird best explained cases of HPAI in premises across Great Britain. Across three distinct epidemic periods between October 2021 and January 2023, 103 species had a significant association in at least one period, and several groups of ecologically similar species were also found to be significant. There was broad consistency between epidemic wave periods (EPs 1 and 3), with groups such as *wildfowl* having consistent significant group-level positive associations with infected premises, alongside some inter-epidemic changes at both the group and species-specific level. In contrast, during the summer period where infected premises were uncommon, our models performed comparatively poorly, failed to find consistent effects across similar species, and wild bird abundance distributions had a less clear association with HPAI occurrence in premises.

Significant positive associations were comparatively much more frequent than negative associations, and consistency between effect sizes of associations across similar species that share morphological and behavioural traits (i.e., group-level effects explored here) helps support a potential causative relationship in these cases. Importantly, despite our method not relying on wild bird HPAIV surveillance or any prior assumptions for susceptibility, many of the associations we have demonstrated are commensurate with our existing understanding of which avian species play a greater role in the dissemination of HPAIVs. Although more than 220 wild bird species have tested positive for HPAIV globally since 2020 (Empres-i, 2023), the primary wild bird species typically associated with maintenance and spread belong to the orders Anseriformes (i.e., wildfowl – ducks, geese, swans) and Charadriiformes (i.e., waders, gulls, terns), with other less commonly implicated orders usually also associated with aquatic habitats such as Gaviiformes (Divers) and Gruiformes (i.e., Rails & Grebes) (Poulson and Brown 2020). However, large-scale, systematic studies of HPAIV are largely lacking in other Orders, despite the limited studies that have been undertaken finding widespread prevalence (albeit often at low levels) (Wille et al. 2023). In this study, *waterbirds* were consistently found as significant predictors in both epidemic waves (EPs 1 and 3) with little change in effect sizes.

We also found that the group *seabirds* showed significant increases in effect sizes during EP3. This matches a widely noted step-change in host-range during the summer of 2022 across Europe, where major HPAI outbreaks were recorded in many seabird species at a scale previously not seen (Caliendo 2023, EFSA 2023B, Pearce-Higgins et al. 2023). Our results for *seabirds* are however largely driven by *gulls & terns* that have more extensive terrestrial distributions than many other seabird species and can often be found terrestrially in the UK year-round. Many other species of *seabirds* across the wider group may therefore be unlikely to have such correlation with outbreaks in captive avian premises. One subgroup we were able to assess and did find significant associations with was *divers*, which was unexpected due to their relatively sparse and coastal distribution in much of the UK and life history that would mean direct contact with captive avian premises being unlikely. However, divers have been shown to have seroprevalence for AIV in other regions, albeit at low levels (Ucher-Koch et al. 2019). They may therefore play a small role in some regions across the coarse spatial scale we assess, with other bridging species driving incursions. Subgroups such as *gulls* are likely to interact with other seabirds, such as Northern Gannets or divers, through the breeding season and may therefore play a key role as a bridge species between HPAI outbreaks in isolated seabird colonies and terrestrial bird populations.

Many of the most conspicuous mass-mortality events within seabirds occurred within species that we could not assess here due to their omission from the eBird modelled species abundance and distribution dataset. This included species such as Northern Gannet and Great Skua where significant proportions of breeding populations were lost (Banyard et al. 2022, Lane et al. 2023). However, these species tend to be present in the UK only in the breeding season and are highly restricted to small islands and coastal edge when not at sea. They are therefore unlikely to have abundance distributions that correlate with HPAIV outbreaks in premises.

We also found significant group-level associations of HPAI cases with abundance of some passerine groupings, such as *sparrows* during EPs 1 and 3. While there is little existing evidence of passerines having a major role as a maintenance reservoir host or bridge species for facilitating premise incursions, this finding may reflect the importance of passerines as an under-studied host species. Furthermore, while passerines tend to show low test positivity rates for HPAI infection (Wade et al. 2023, EFSA 2022) and may therefore have limited capacity as biological sources of virus (i.e., virus replicates inside the wild host), they may also play a role as mechanical transporters via fomite (e.g., virus carried externally on feet or feathers). External swabbing of small passerines in the United States during previous outbreaks found no support for mechanical transport potential via fomites (Houston et al. 2017). However, small sample sizes together with large populations typical of small passerines lead to low confidence in prevalence. If passerines are acting as sources of transmission (mechanical or biological), many species are small and can be difficult to exclude from poultry premises or adjacent habitats via biosecurity measures unless bird exclusion measures are of high standard and well-maintained, and therefore may pose a significant ongoing risk of incursion. Traditional surveillance programs may also be likely to miss mechanical transmission of virus in these species, as testing is often targeted towards detection of active or past infection.

Of particular note was the association of HPAI infected premises with free-living *non-native gamebirds* during EP 3. Although exact numbers are undefined, tens of millions of non-native gamebirds are reared and released in the United Kingdom each year to bolster feral breeding populations (breeding pop. est. ∼4.4 million; Blackburn and Gaston 2018) for the purpose of gamebird shooting (Madden 2021). Both Common Pheasant *Phasianus colchicus* and Red-legged Partridge *Alectoris rufa*, the two non-native gamebirds released in highest numbers annually, had species-specific positive associations during EP 3. EBird-modelled population abundances for non-native gamebirds are not a direct measure of gamebird release locations. However, it is expected that there would be a high degree of correlation between the two as released birds typically only disperse a short distance from release sites and have low survival rates following release. For instance, typically 90–95% of released pheasants and partridges remain within ∼1 km of release sites and only ∼15% of releases survive to the end of the shooting season (Madden et al. 2018). Significant numbers of Mallard *Anas platyrhynchos* are also released for shooting within this period (estimated ∼2.6 million per year compared to ∼31.5 and ∼9.1 million Common Pheasant and Red-legged Partridge, respectively; Madden 2021). However, while Mallard also had a significant positive association during EP 3, it was not possible to consider truly wild and released birds separately.

The role of reared and released birds in HPAI is controversial; in 2022 the Royal Society for the Protection of Birds (RSPB) called for a moratorium on gamebird releases as a precautionary measure to reduce the spread of HPAI in wild birds, an approach dismissed by the British Association for Shooting and Conservation (BASC) citing a lack of supporting evidence (Gray and Loeb 2022). Despite cases of HPAI within gamebird hatcheries and rearing premises (Fujiwara et al. 2022; DEFRA 2023A), there remains a lack of conclusive evidence for the role of released gamebirds in the spread of virus to poultry or wild birds. The likelihood of incursions into the gamebird sector was considered low based on egg and chick movements (Fujiwara et al. 2022) but higher during “catching-up” (when surviving released gamebirds may be re-caught after cessation of the shooting period; DEFRA 2023B). However, the risk of HPAI spread from free-roaming gamebirds to wild birds post-release was deemed high and has potential to contribute to viral maintenance in other wild bird populations, ultimately leading to increased infection pressure (DEFRA 2022). Furthermore, experimental *in vivo* inoculation with clade 2.3.4.4b H5N6 has shown that Common Pheasant is capable of both acquiring HPAIV and subsequently transmitting to poultry (Liang et al. 2022). Similar outcomes were seen following experimental infection of pheasants with clade 2.3.4.4b H5N1 and H5N8 viruses, with pheasants being more susceptible to infection than red legged partridges (Seekings et al., 2023). The significant positive associations with *non-native gamebirds* that we identify here strongly suggest that increasing surveillance of this group is necessary to more robustly assess whether non-native gamebirds are a key source of HPAI spillover onto captive avian premises, particularly following release.

Spatial associations between wild bird abundance and HPAI cases in premises provide tentative evidence of which species may be driving viral incursions (either directly onto premises or indirectly by acting as local amplifiers), with a higher species abundance increasing the likelihood of cases in that area. However, purely associative relationships such as these cannot be used as unequivocal evidence of causation. Associations may arise from chance alone, or due to co-linear effects rather than a causal link. Indeed, this is likely to be the driver for the few negative associations found in this study, as there is no clear causative hypothesis for higher abundance of a wild bird species driving lower HPAI incidence in premises. It is perhaps more likely for many species with negative associations that their abundance is either negatively correlated with species that are drivers of HPAI incursions or they arose by random chance.

Despite these limitations, our approach benefits from the ability for rapid deployment without the requirement for extensive prior surveillance of disease in all potential host species, which is often financially and logistically prohibitive. Even in developed countries such as Great Britain, logistical constraints such as the reporting, triage, collection, and transport of carcasses, as well as legal restrictions on sample handling, can limit testing capacity. Notably, we saw a marked decline in positive HPAI tests in wild birds across our three EPs despite an increase in premise cases (S6 Fig). Indeed, insufficient positive wild bird cases may be the cause for the lack of predictive power observed in EP 3, where wild bird HPAI cases did not significantly predict premise cases.

Our research shows that the use of publicly available wild bird abundance and distribution data can complement surveillance in wild birds to help identify species that should be prioritised for testing in close to real time. This could help to target limited surveillance resources and to monitor potential changes in infected wild species even in the absence of high mortality. Rapid deployment may also aid in risk mitigation for incursions onto premises, by enabling improved biosecurity measures to be put in place with minimal delay that are better tailored towards higher-risk species. The use of eBird data is particularly valuable because predictions are made at the global scale, at relatively high resolution, and are free to access for over 2000 species (Strimas-Mackey et al. 2022). However, continued efforts may be needed to improve capacity building in regions where the requisite citizen science data may be lacking, and species abundance distributions cannot be accurately modelled.

Despite our analyses largely relying on publicly available data, we controlled for avian livestock and farm density using poultry distribution datasets that are not publicly available in Great Britain. Controlling for avian livestock density may be somewhat achievable globally with The Gridded Livestock of the World 3 (Gilbert et al. 2018; ∼10km^2^ resolution), but farm density data may only be available for some countries. In our analyses, we used The Great British Poultry Register (GBPR), as it enabled poultry stock and farm density to be calculated at much finer spatial resolutions, yet it is not publicly available due to data protection constraints and is only compulsory for sites with >50 birds. We also lack sufficient data on how biosecurity (in terms of preventing wild bird incursion) may vary across premises, either by type or size, and whether there are significant spatial patterns in this across Great Britain. Improved understanding of these potential biases would enable more accurate spatial associations between wild bird abundance and HPAI case risk in premises.

## 5. Conclusions

Publicly available wild bird abundance and distribution predictions developed using citizen science bird-counting initiatives, such as those offered by eBird, can be a powerful tool in helping to identify potential drivers of wild-bird mediated HPAIV incursions into premises. Higher abundance in avian groups such as wildfowl (ducks, geese, etc.) was found to be consistently associated with a higher incidence of HPAI cases in premises between epidemic waves. Some avian species groups became more important in the most recent epidemic wave, perhaps linked to a change in viral host range, or species-specific drivers such as *non-native gamebirds* only being associated with HPAI in premises during epidemic waves coinciding with mass-release. These associations may help guide future targeted mass-surveillance and aid understanding of the changing host-range of HPAIV as it continues to adapt and spread in wild bird populations.

## Author statements

### Ethical approval

Not applicable.

## Acknowledgements

This material uses data from the eBird Status and Trends Project at the Cornell Lab of Ornithology, eBird.org. Any opinions, findings, and conclusions or recommendations expressed in this material are those of the author(s) and do not necessarily reflect the views of the Cornell Lab of Ornithology.

## Supplementary Material

**S1 Table.**
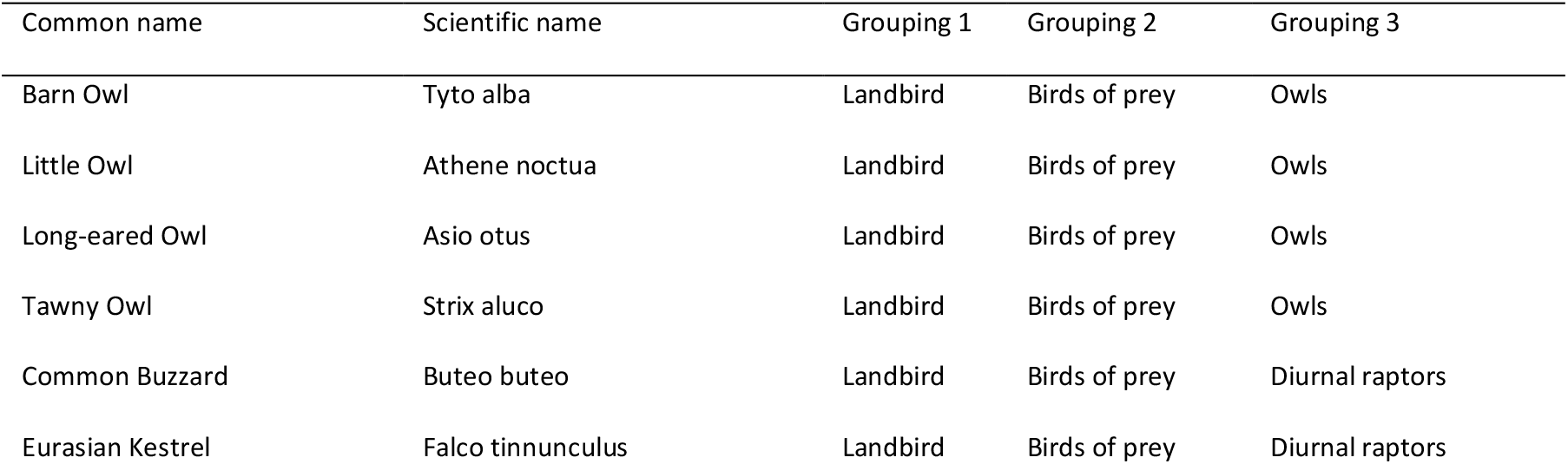
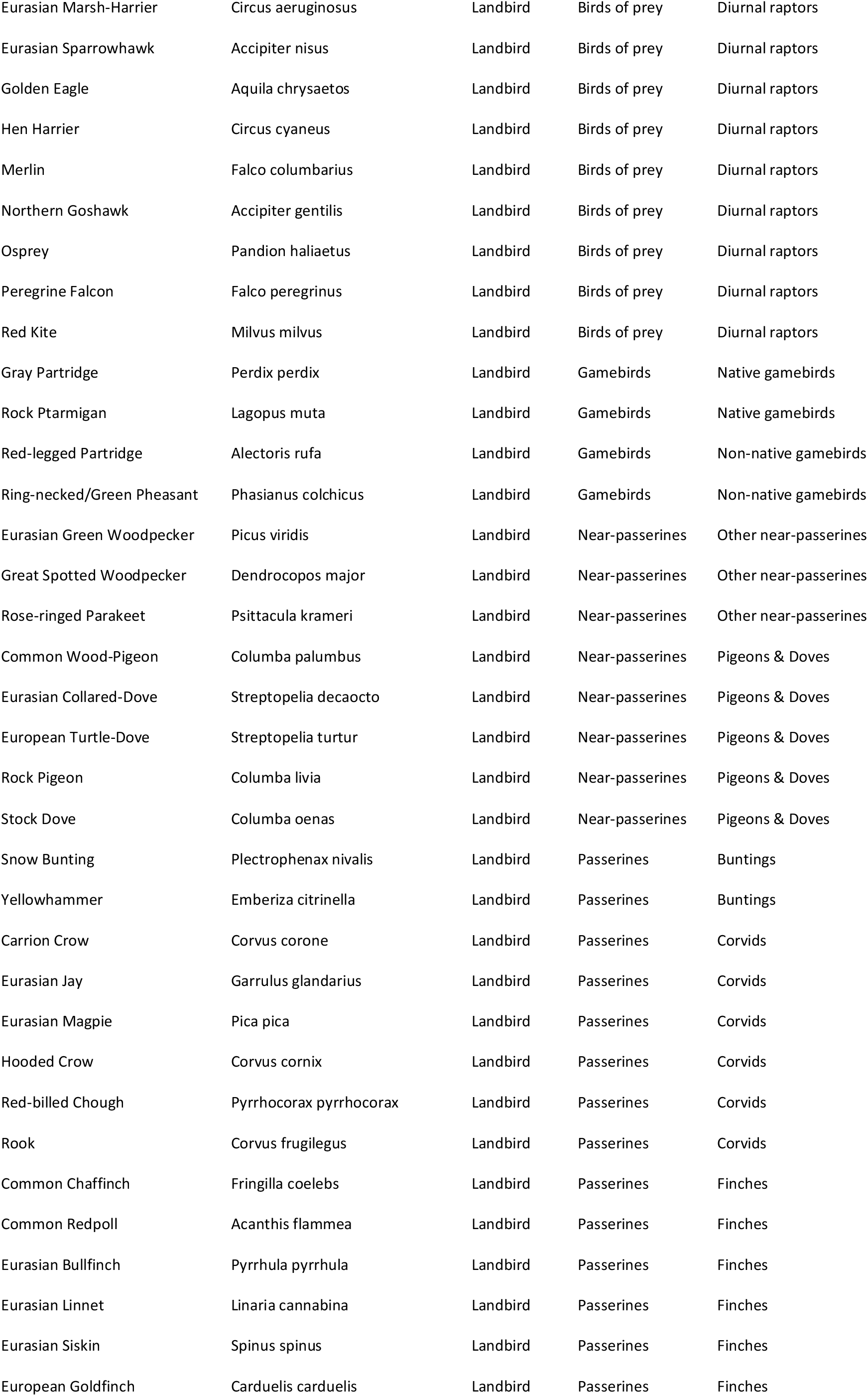
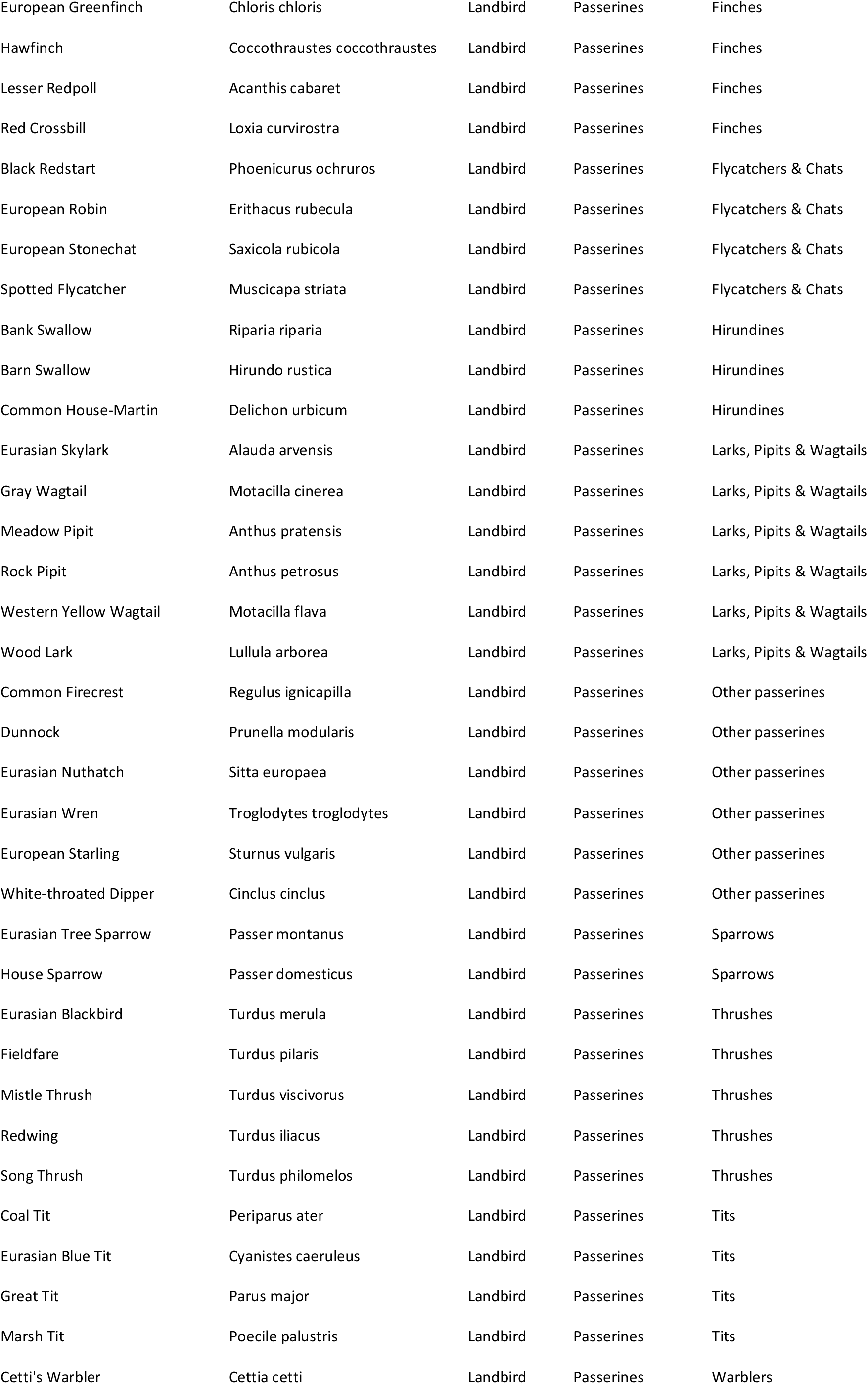
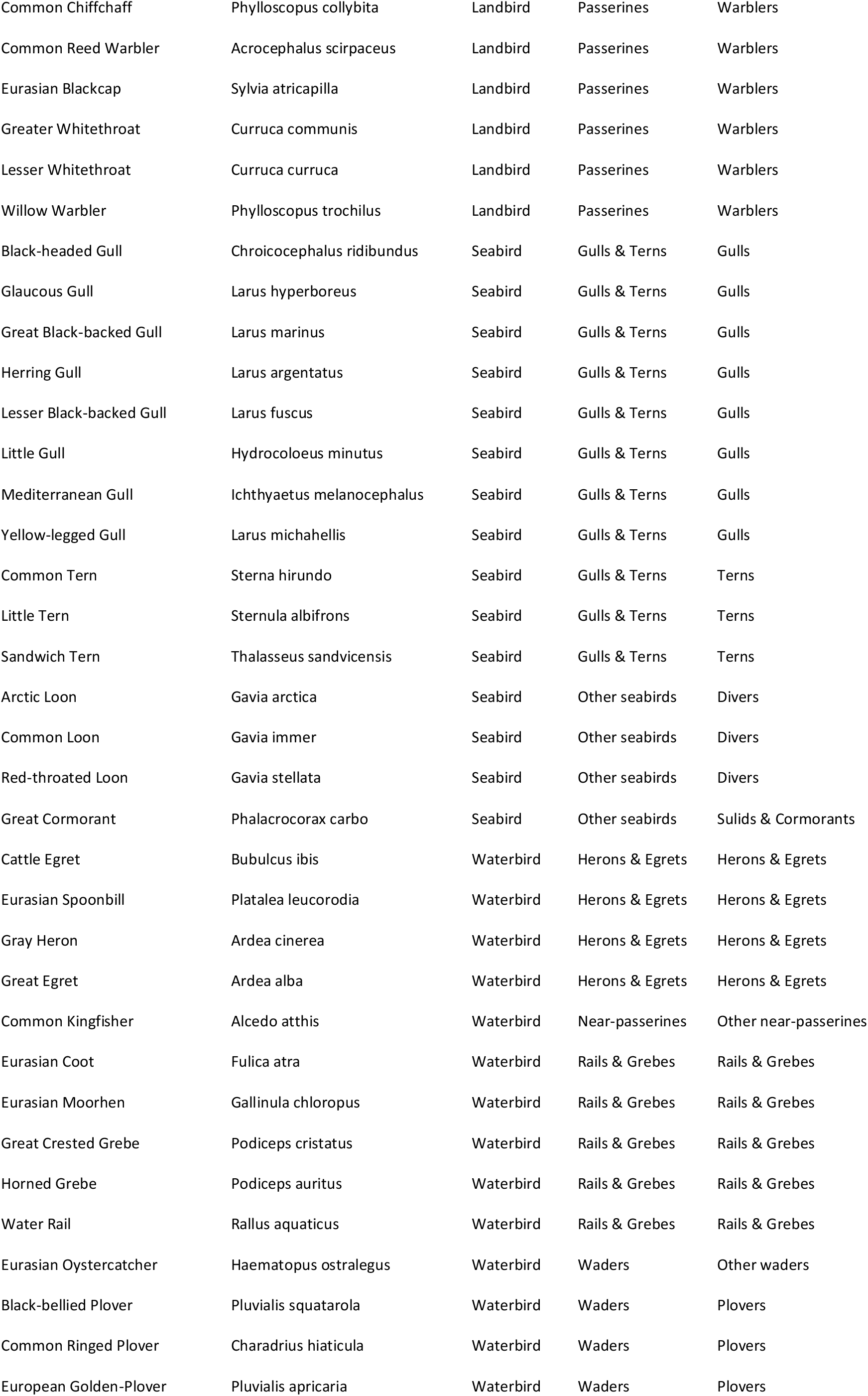
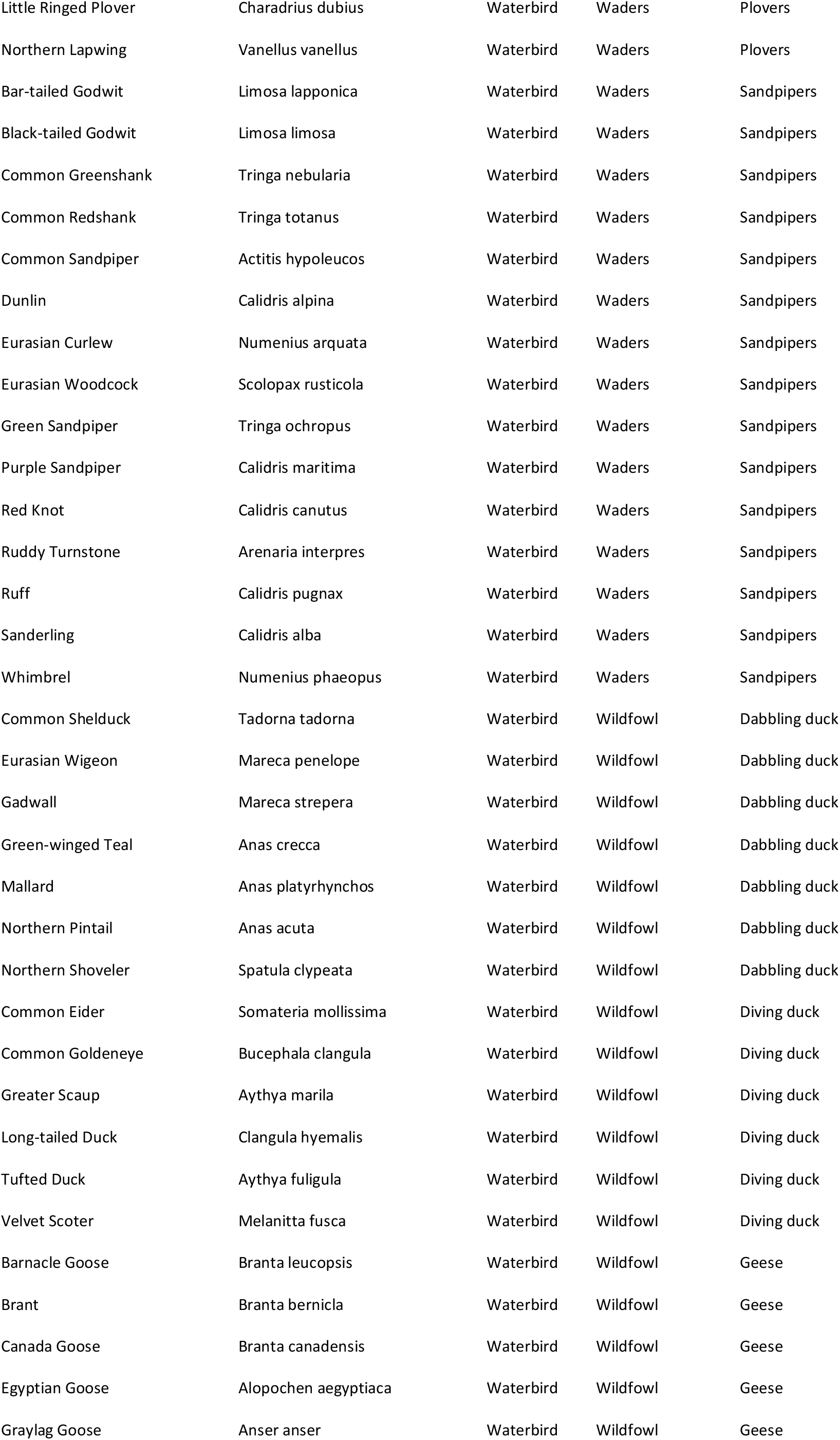
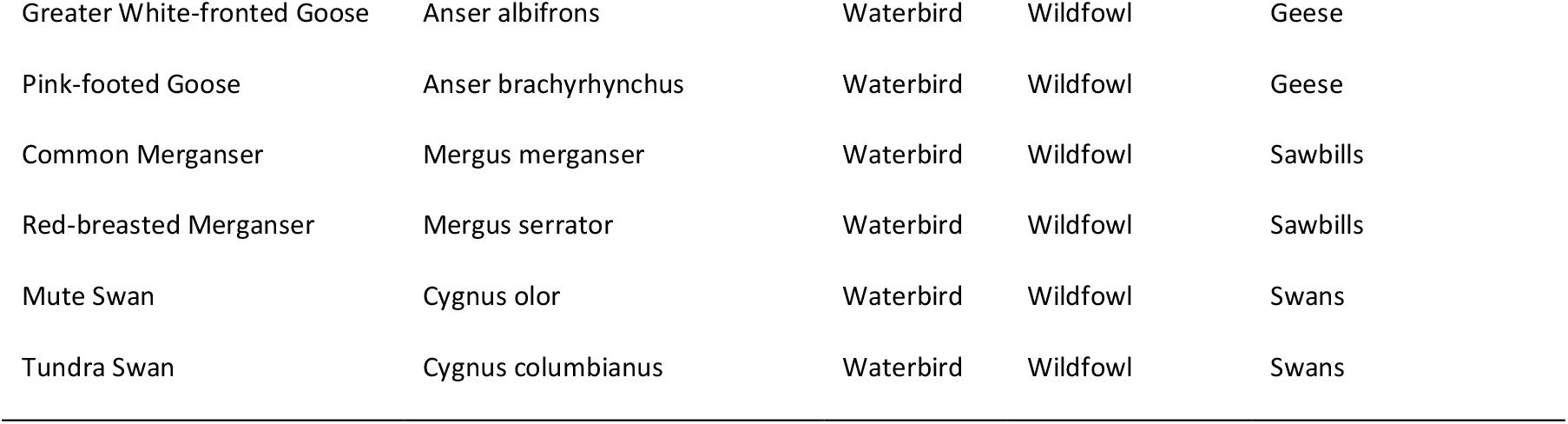
Species groupings

**S2 Table.** Linear model estimated mean slope coefficients across species groupings and epidemic periods. SE = standard error, df = degrees of freedom.

| Epidemic period | Grouping | Estimated mean | SE | df | t ratio | p value |
| --- | --- | --- | --- | --- | --- | --- |
| Epidemic period 1 | Landbird | -0.046 | 0.036 | 374 | -1.286 | 0.199 |
| Epidemic period 2 | Landbird | -0.064 | 0.052 | 374 | -1.235 | 0.218 |
| Epidemic period 3 | Landbird | 0.043 | 0.023 | 374 | 1.886 | 0.060 |
| Epidemic period 1 | Seabird | 0.166 | 0.087 | 374 | 1.907 | 0.057 |
| Epidemic period 2 | Seabird | 0.160 | 0.122 | 374 | 1.307 | 0.192 |
| Epidemic period 3 | Seabird | 0.270 | 0.059 | 374 | 4.541 | **<0.001** |
| Epidemic period 1 | Waterbird | 0.259 | 0.029 | 374 | 8.833 | **<0.001** |
| Epidemic period 2 | Waterbird | -0.070 | 0.065 | 374 | -1.080 | 0.281 |
| Epidemic period 3 | Waterbird | 0.168 | 0.023 | 374 | 7.330 | **<0.001** |

**S3 Table.**
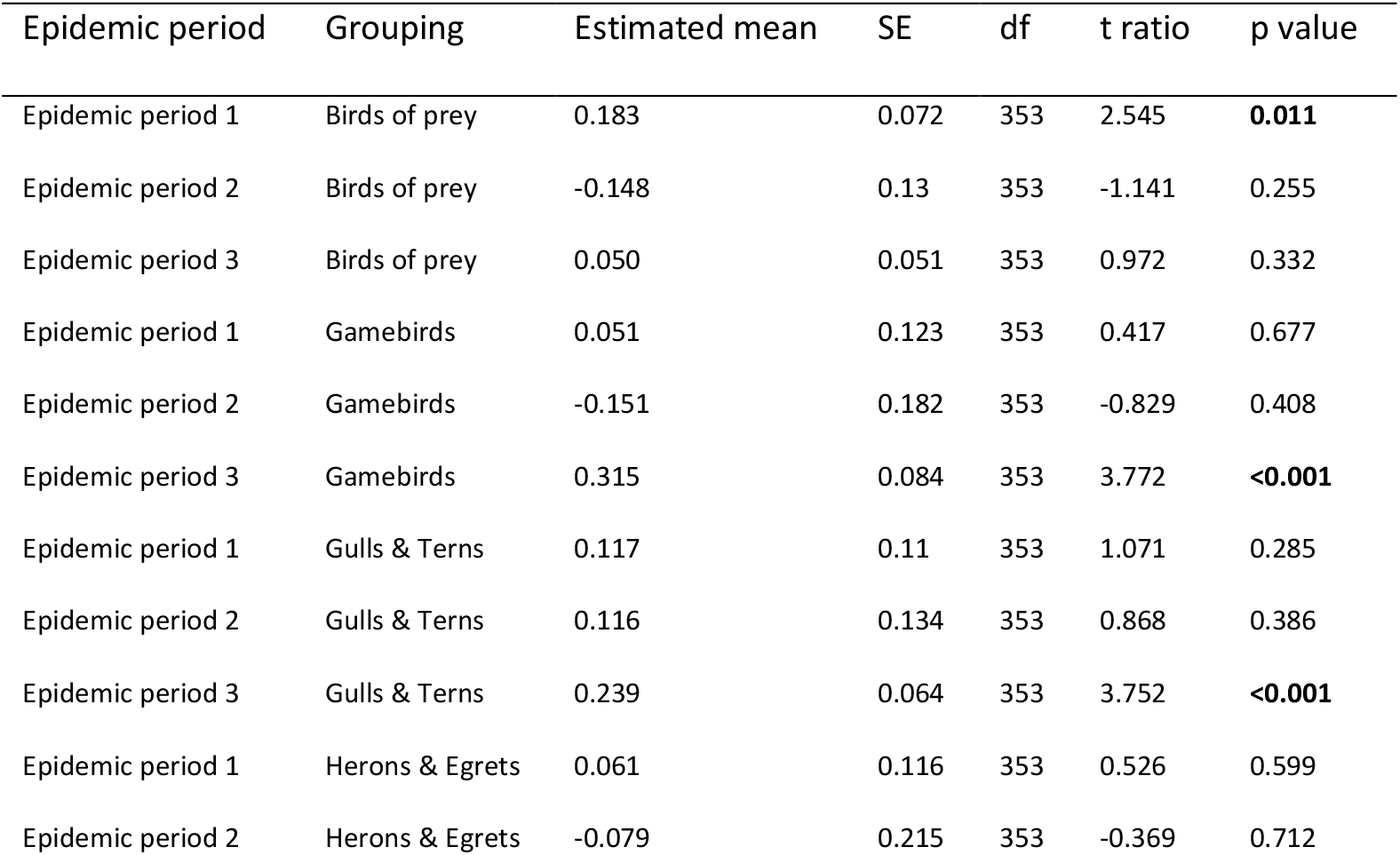
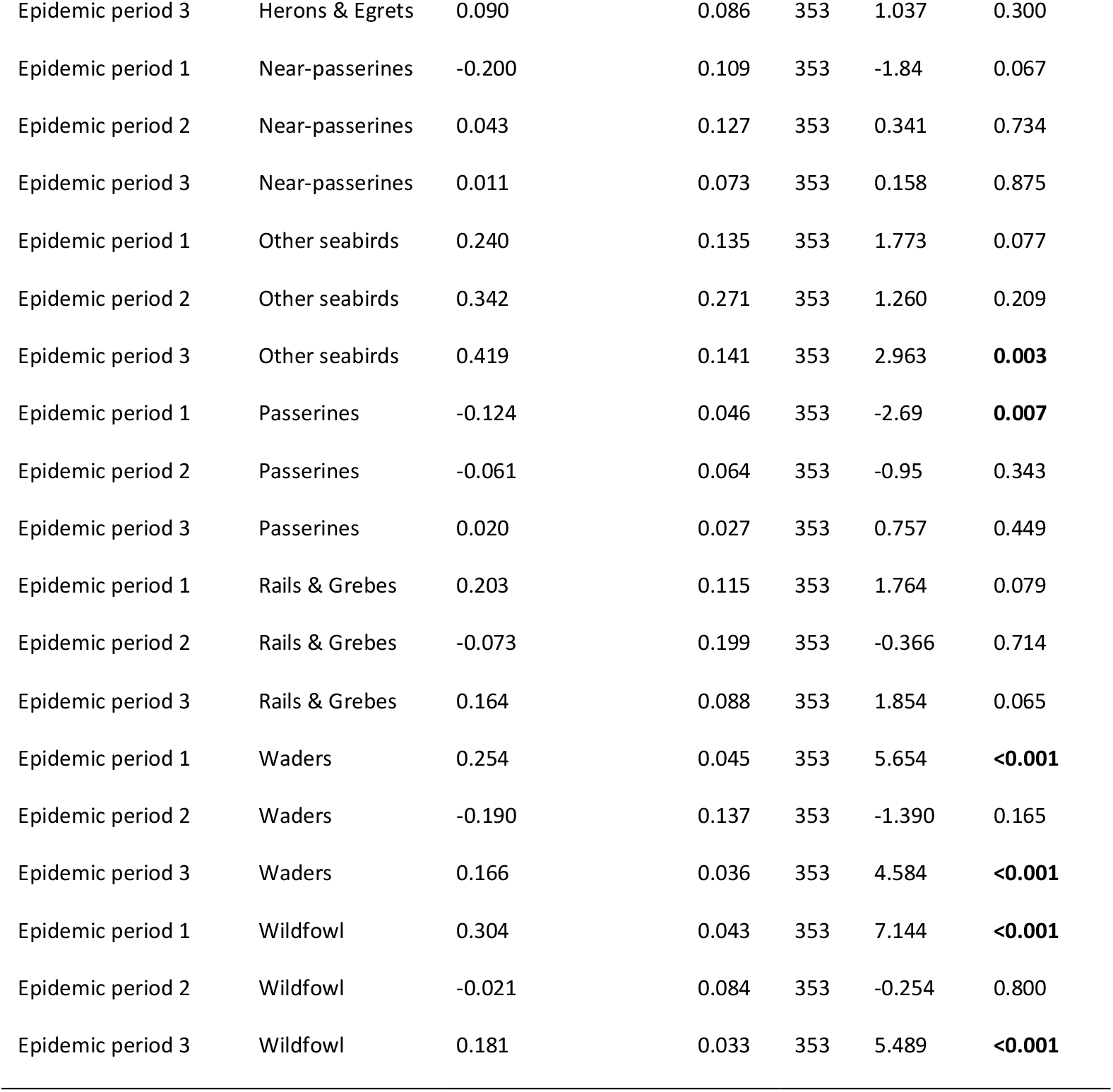
Linear model estimated mean slope coefficients across species groupings and epidemic periods. SE = standard error, df = degrees of freedom.

**S4 Table.**
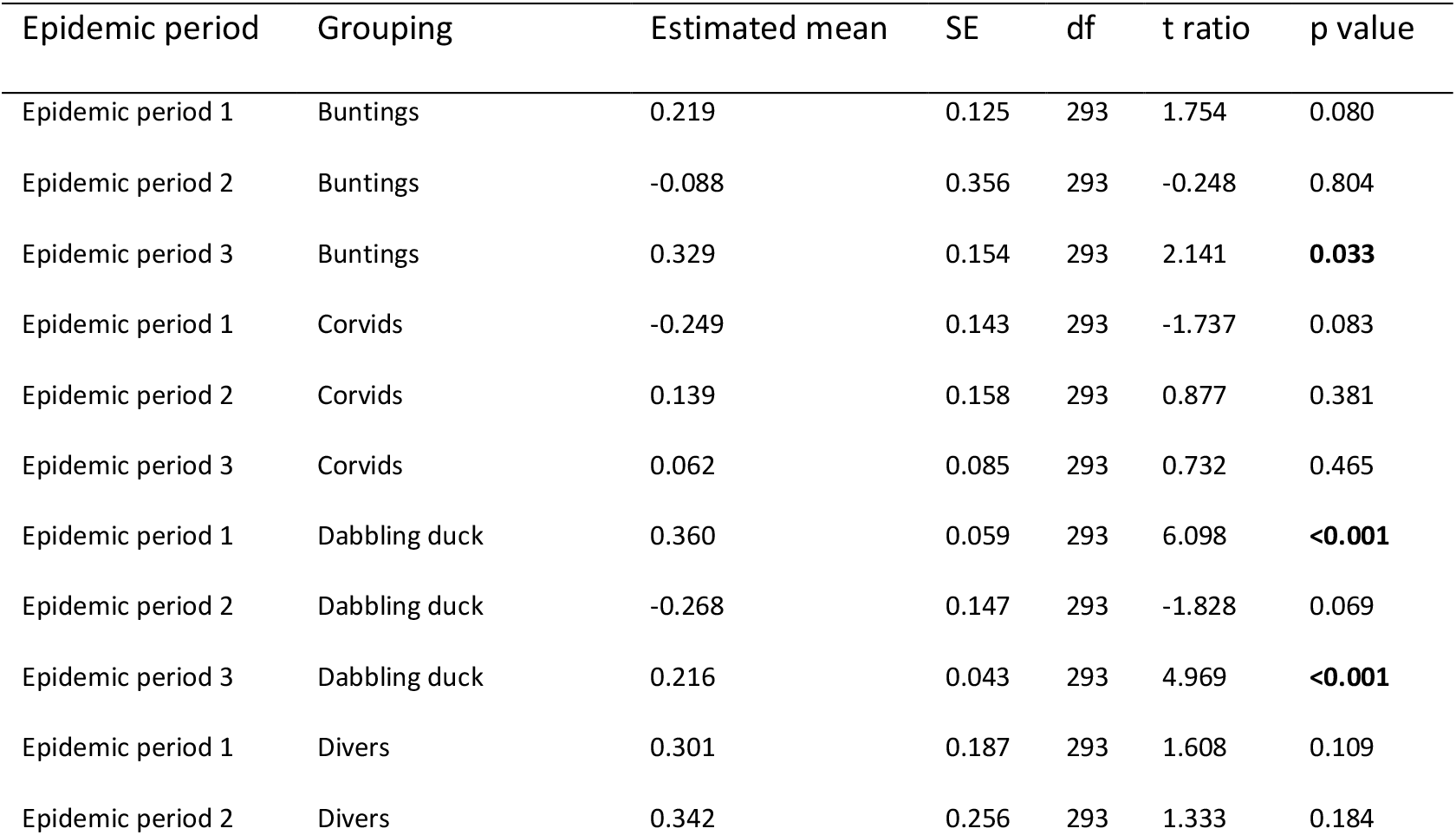
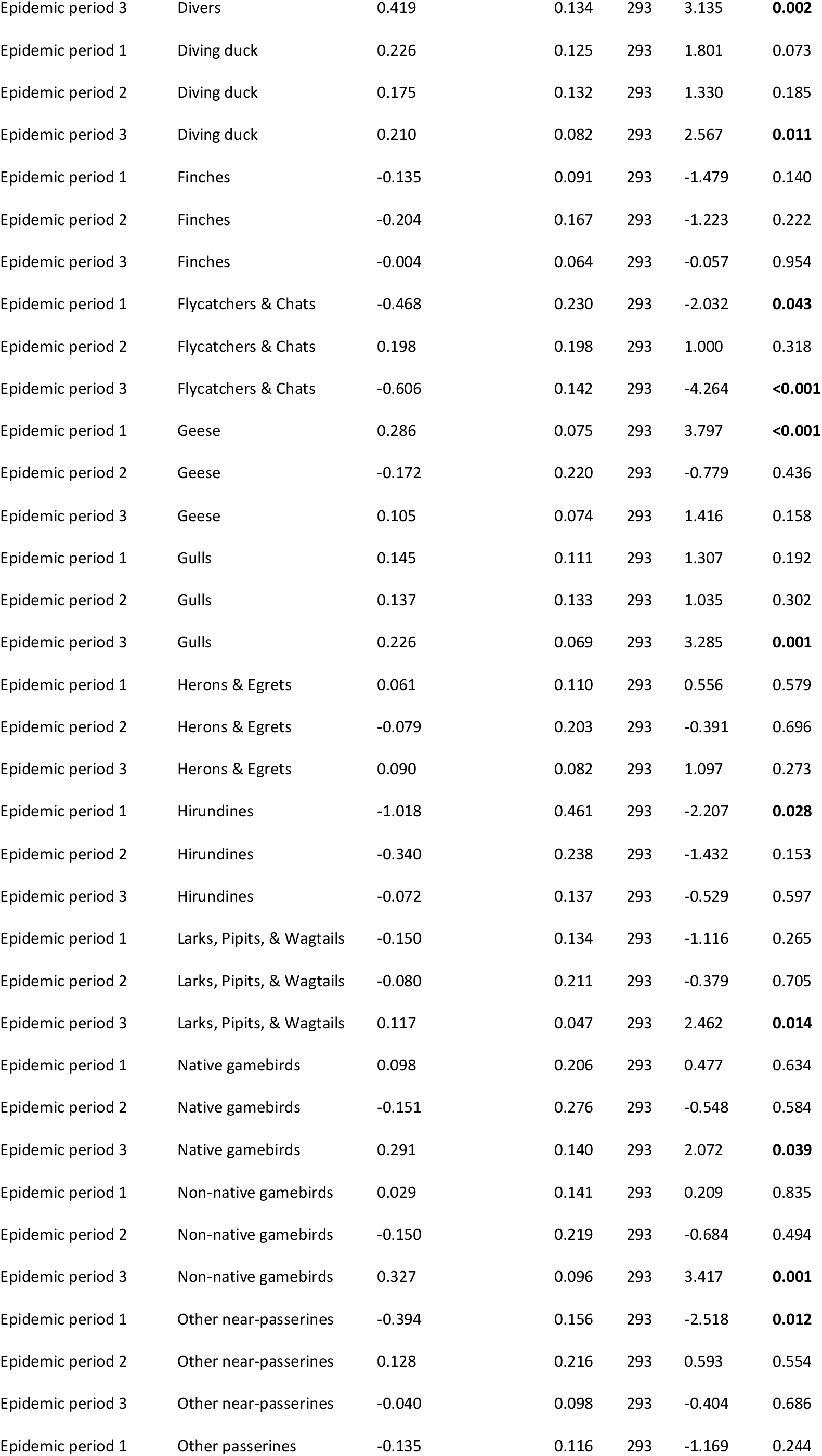
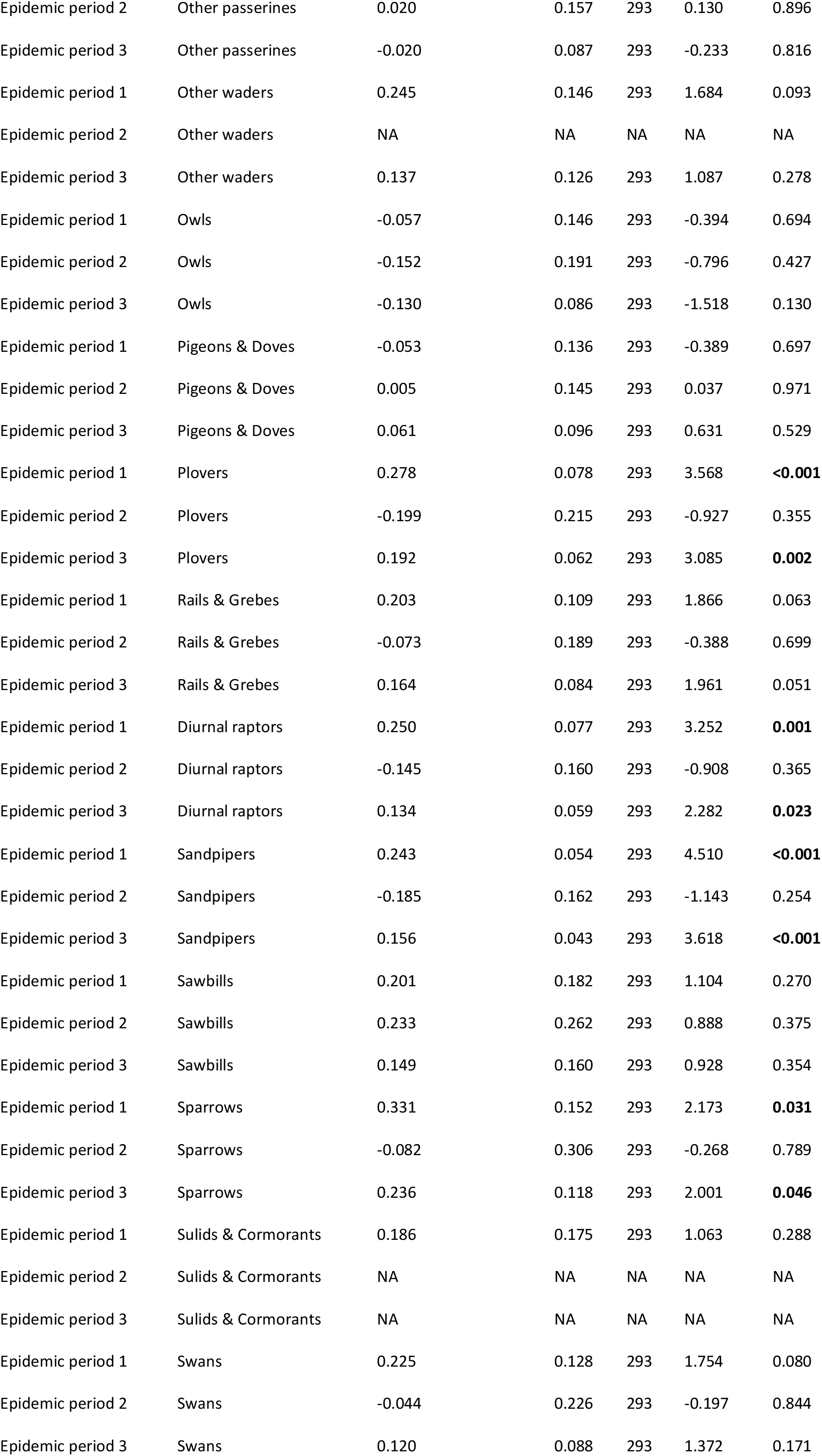
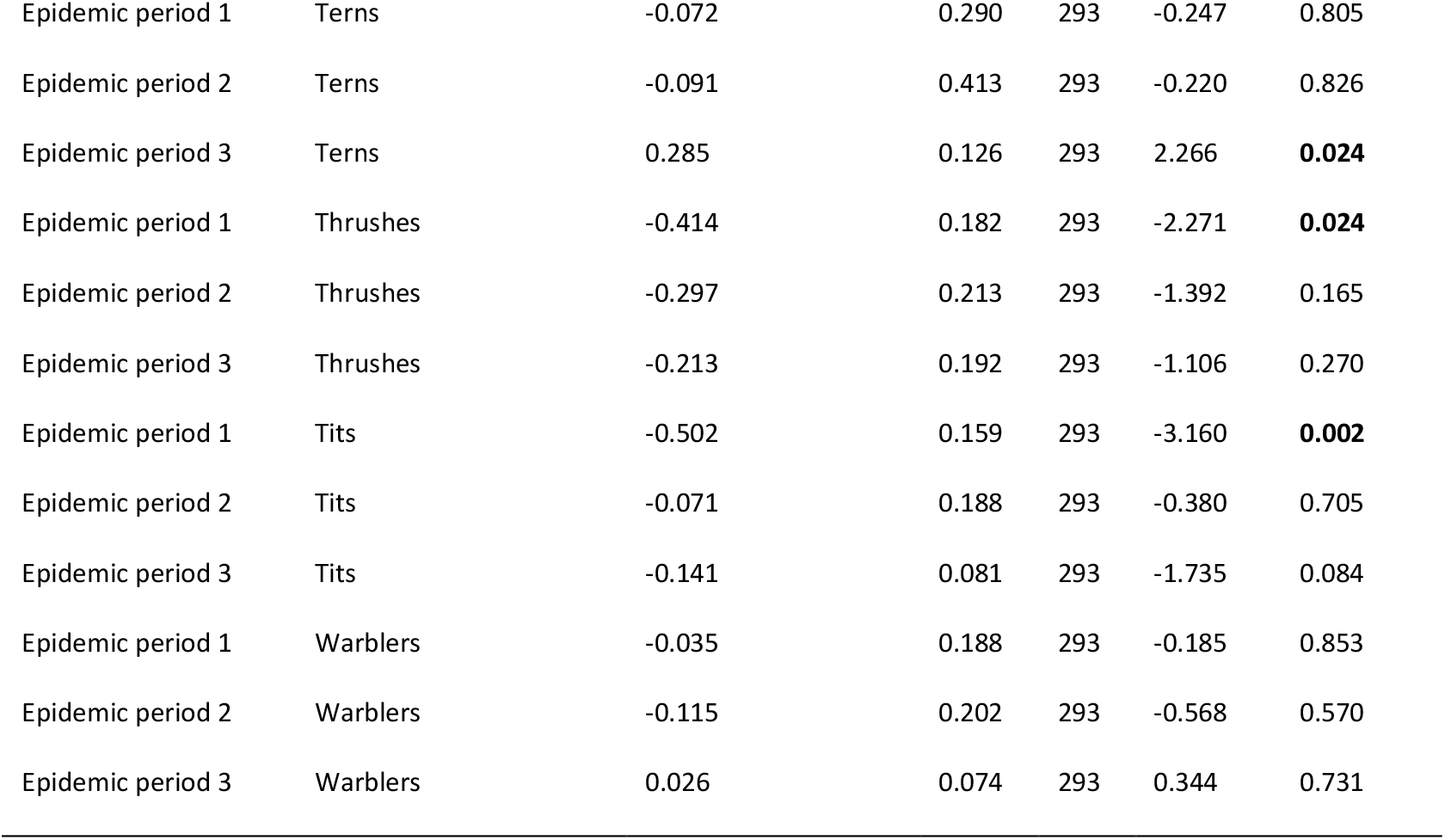
Linear model estimated mean slope coefficients across species groupings and epidemic periods. SE = standard error, df = degrees of freedom.

**S5 Table.**
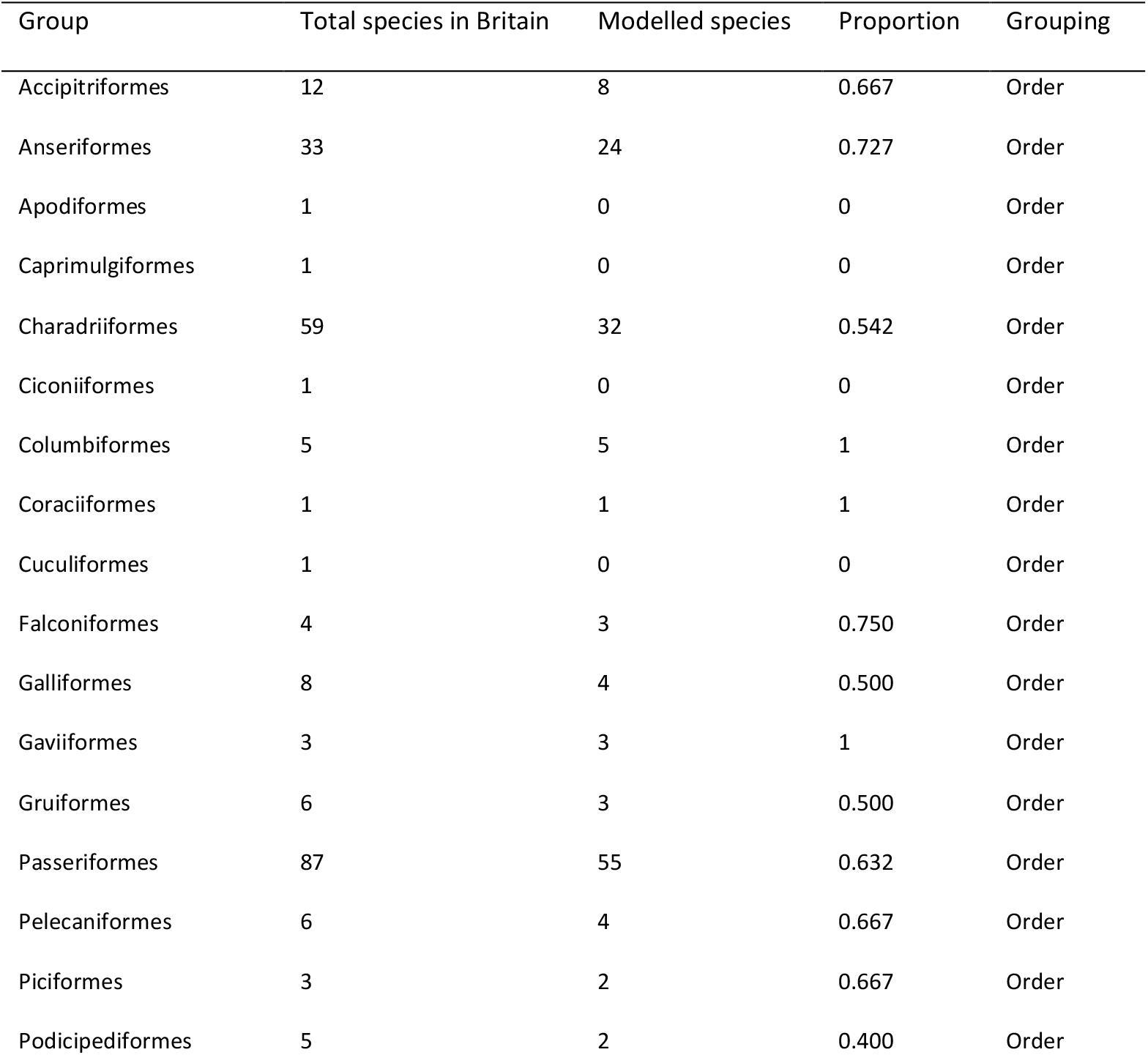
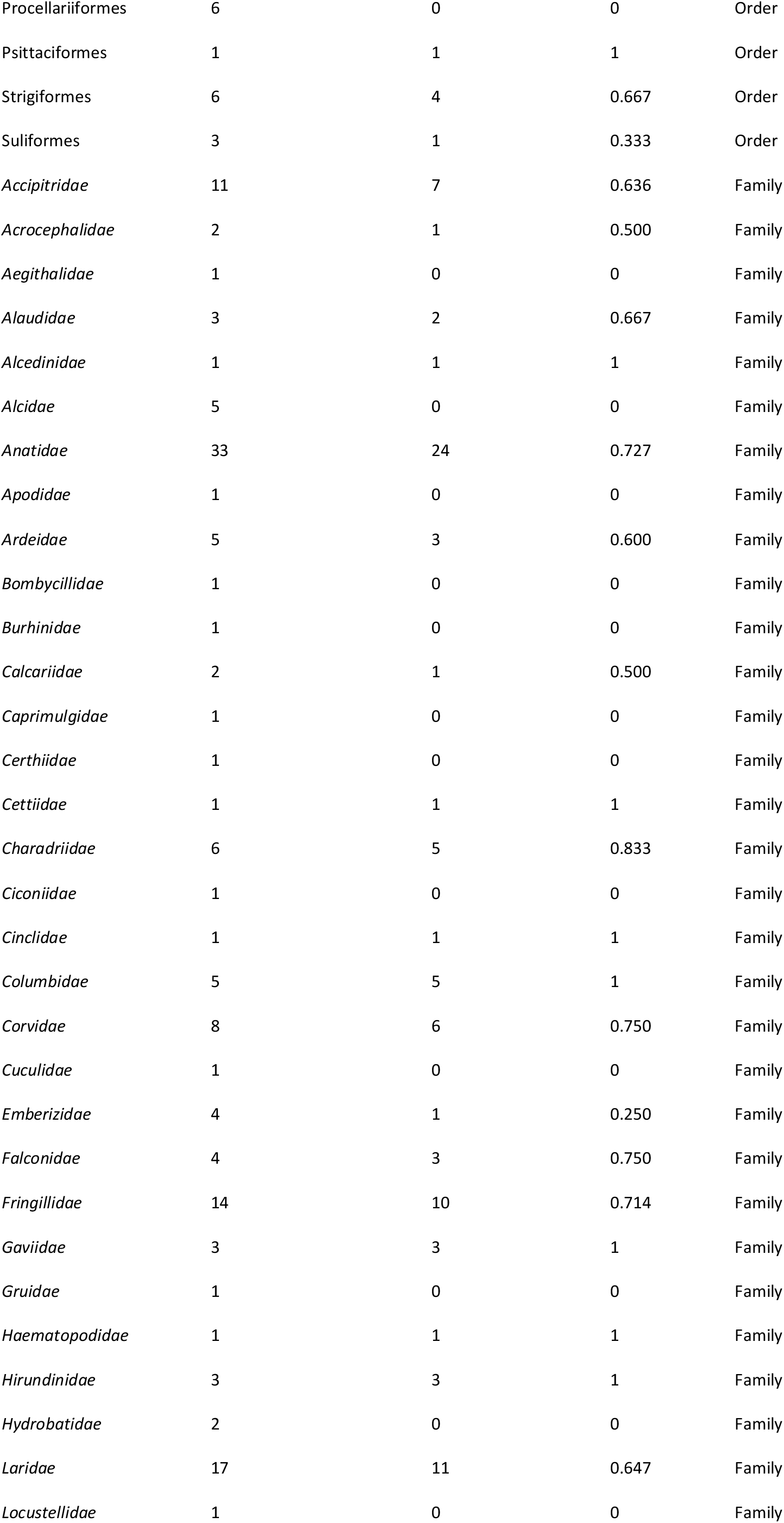
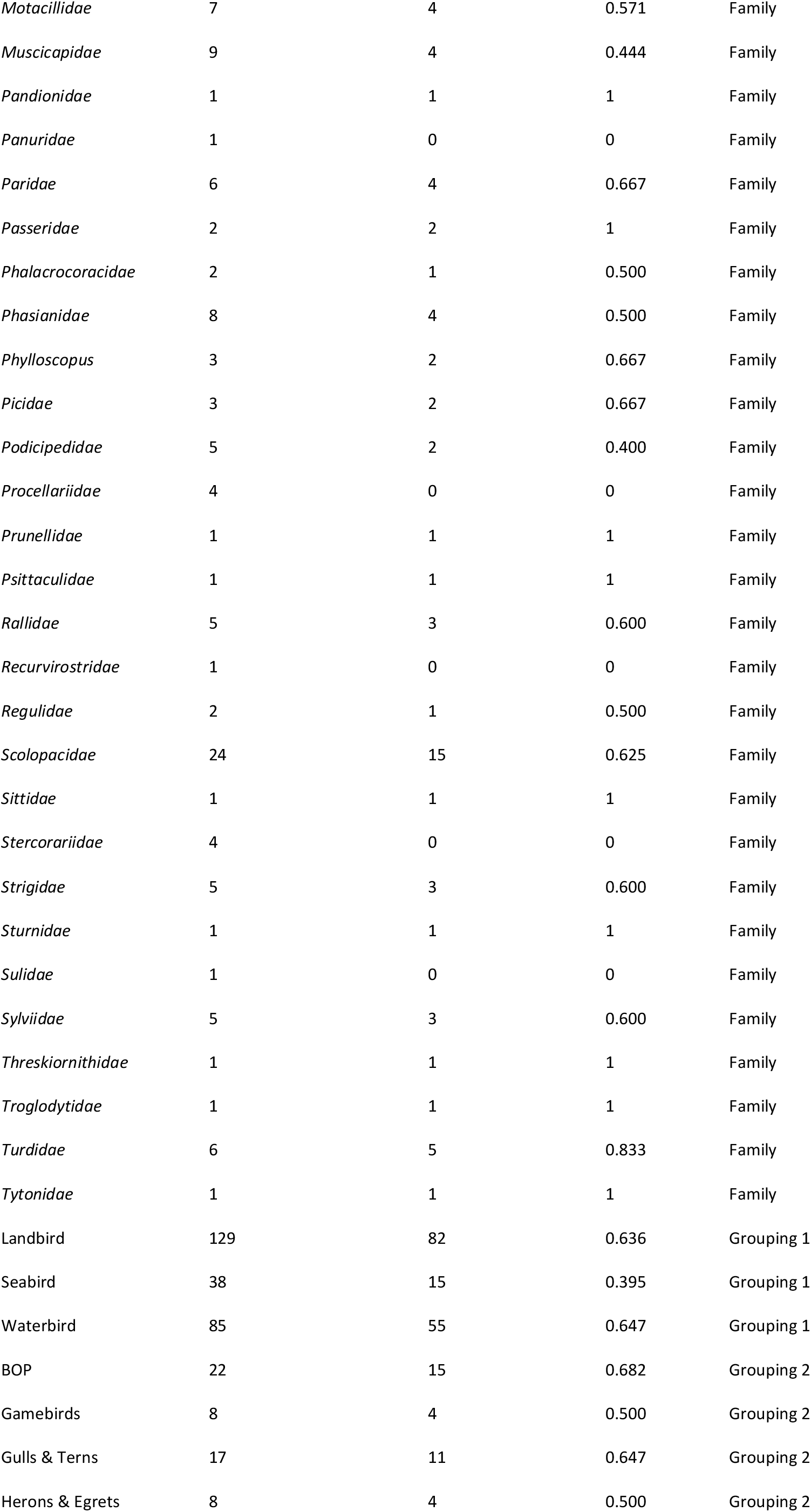
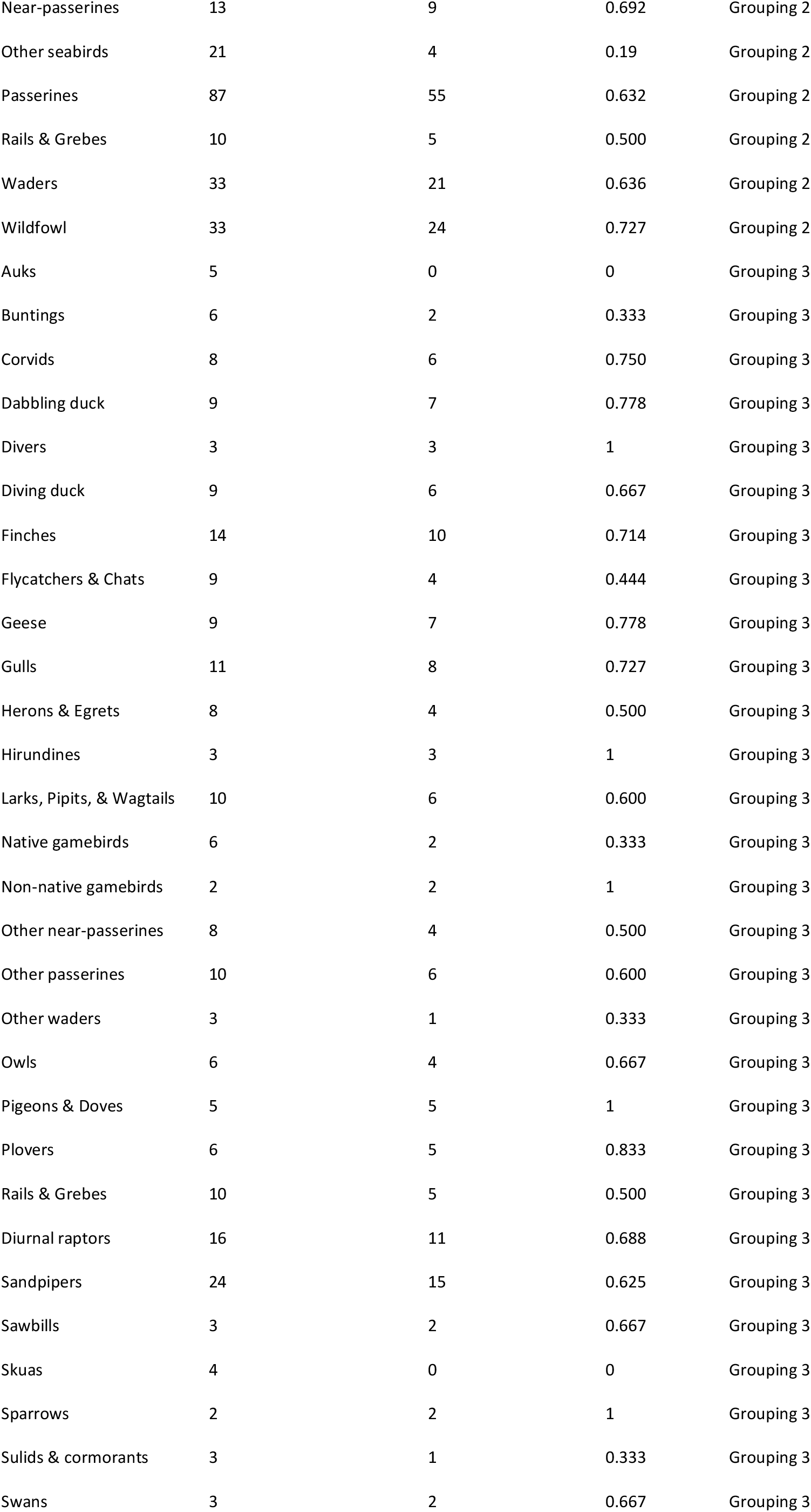
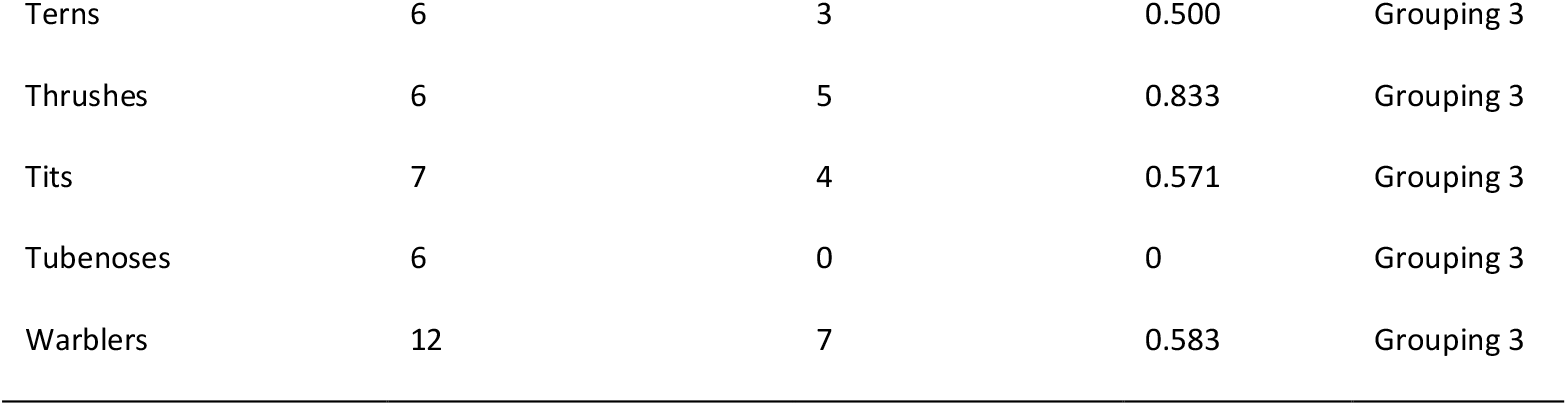
Counts of species within each of our grouping variables and the number of those in which we were able to model associations with HPAI cases in premises. Total counts are based upon the BOU British List excluding vagrants.

**S6 Table.**
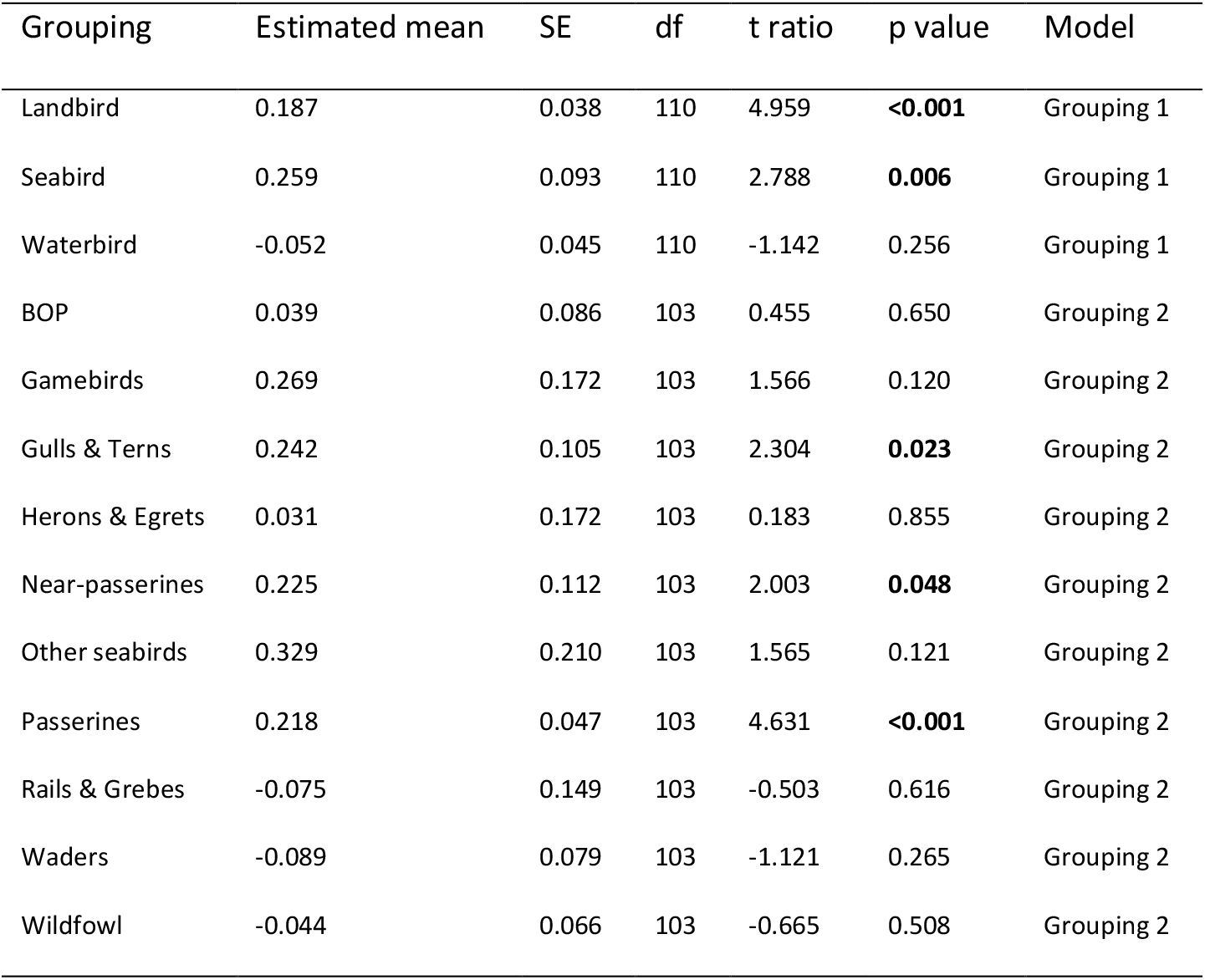
Linear model estimated mean change in species abundance slope coefficients between epidemic period 3 and epidemic period 1, across species groupings. SE = standard error, df = degrees of freedom.

**S1 Fig.**
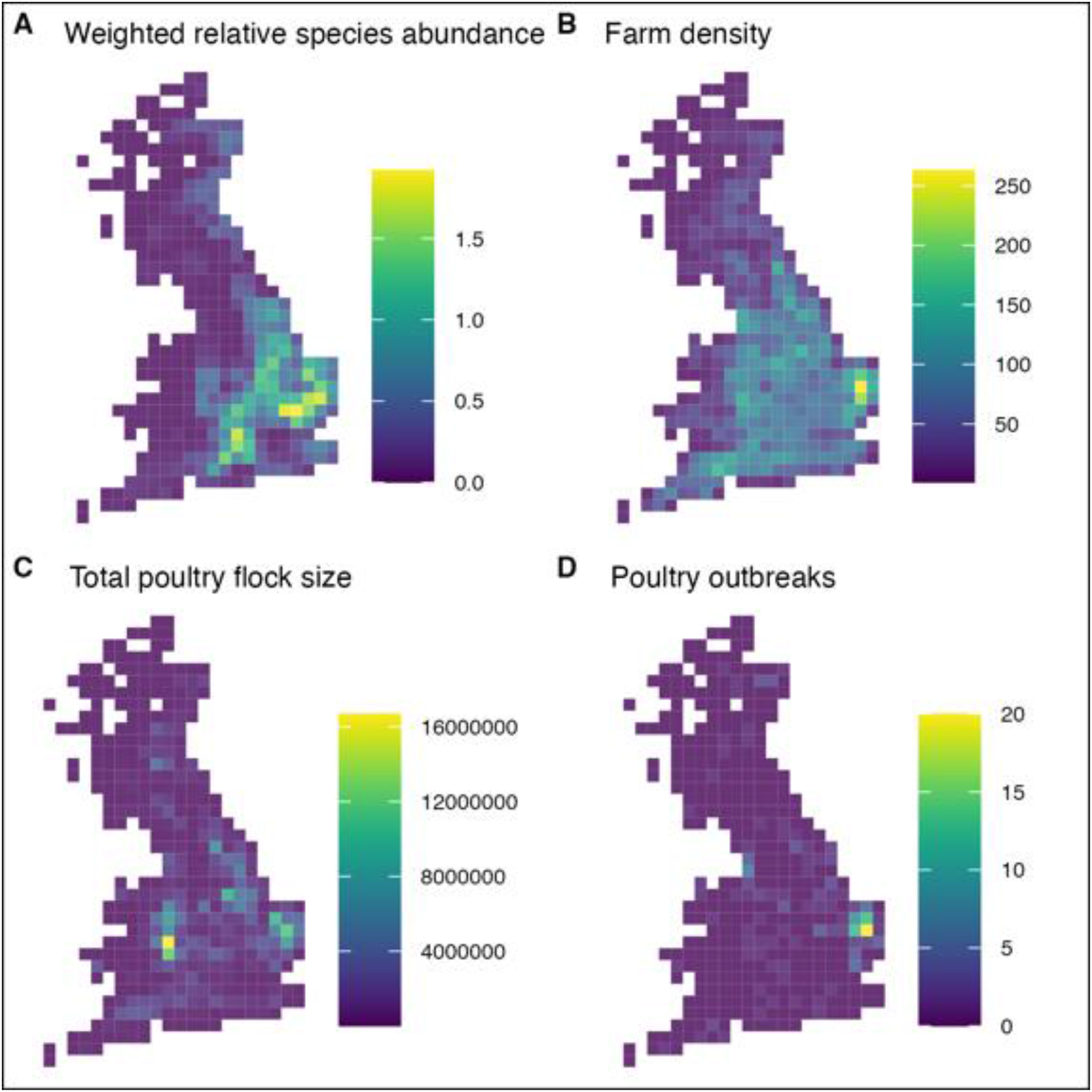
Example spatial data used within our modelling process. Maps are for Yellowhammer Emberiza citrinella within epidemic period 3 (17^th^ August 2022 – 20^th^ January 2023). Each spatial unit is 26.7km^2^. **A)** Weighted sum of relative species abundance derived from eBird Status and Trends species relative abundance modelling weighted by the weekly proportion of HPAI outbreaks in premises within that period. **B)** Count of premises with >50 captive birds within each spatial unit as listed in the Great British Poultry Register. **C)** Total poultry flock size with each spatial unit as listed in the Great British Poultry Register. **D)** Count of HPAI cases in premises within the period.

**S2 Fig.**
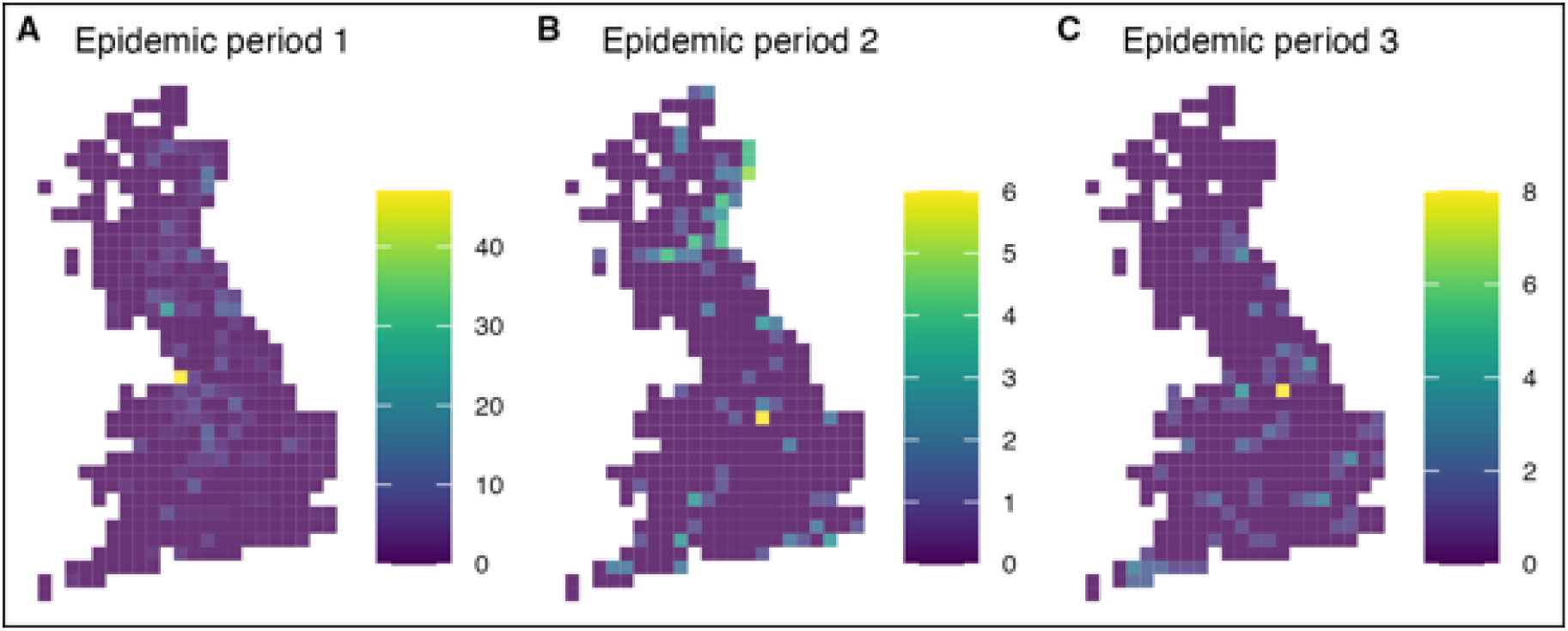
Rasterized counts of positive wild bird HPAI cases from the Empres-i database within the three distinct epidemic periods (EP1 = 19^th^ October 2021 – 9^th^ February 2022; EP2 = 10^th^ February 2022 – 16^th^ August 2022; EP3 = 17^th^ August 2022 – 20^th^ January 2023).

**S3 Fig.**
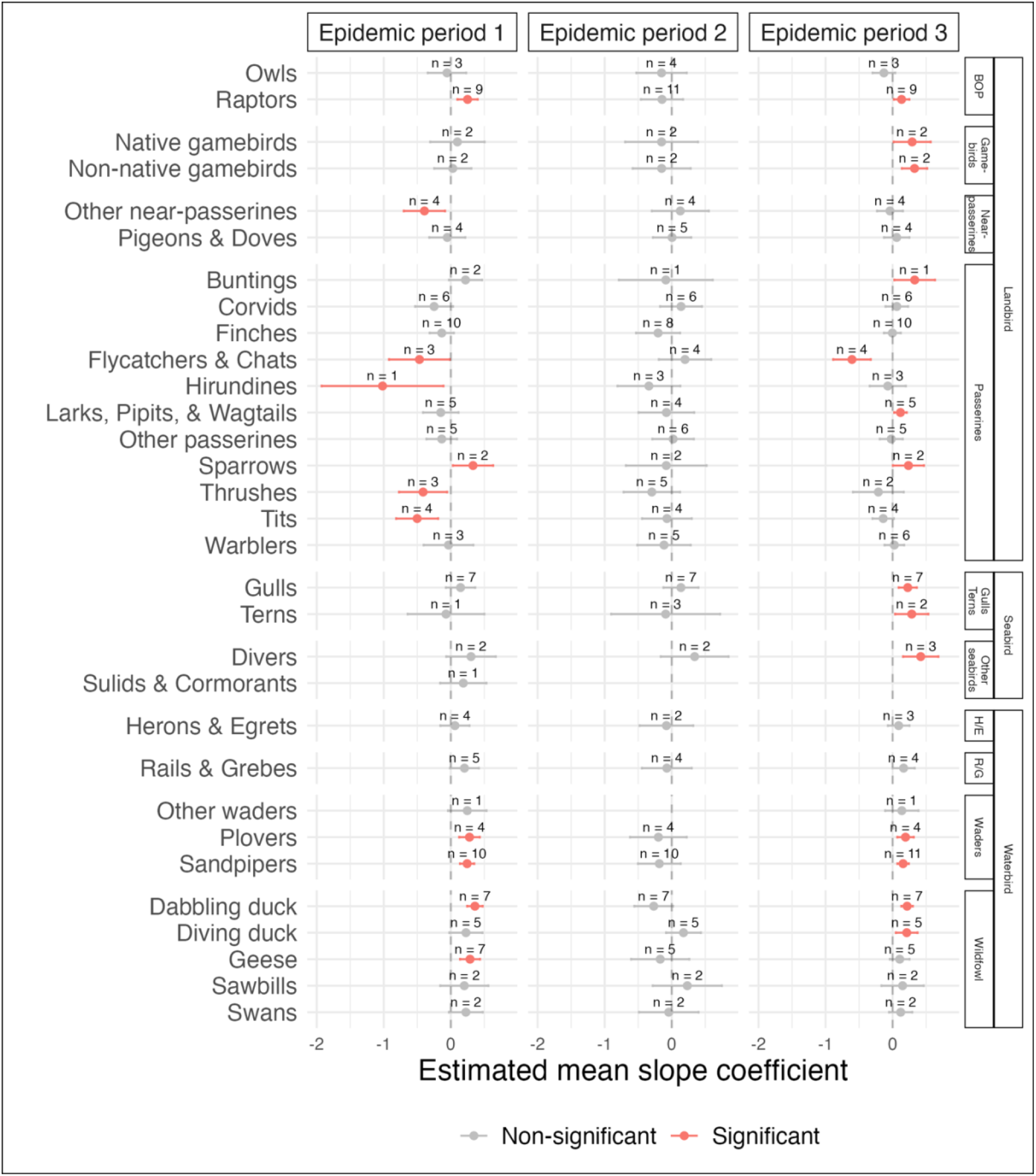
Estimated third-order group-level means for linear model coefficients describing effects of weighted relative species abundance on probability of HPAI cases in premises. Estimated means derived from a GLM assessing group effects across epidemic periods (EP1 = 19^th^ October 2021 – 9^th^ February 2022; EP2 = 10^th^ February 2022 – 16^th^ August 2022; EP3 = 17^th^ August 2022 – 20^th^ January 2023). ‘H/G’ = Herons & Egrets, ‘R/G’ = Rails & Grebes.

**S4 Fig.**
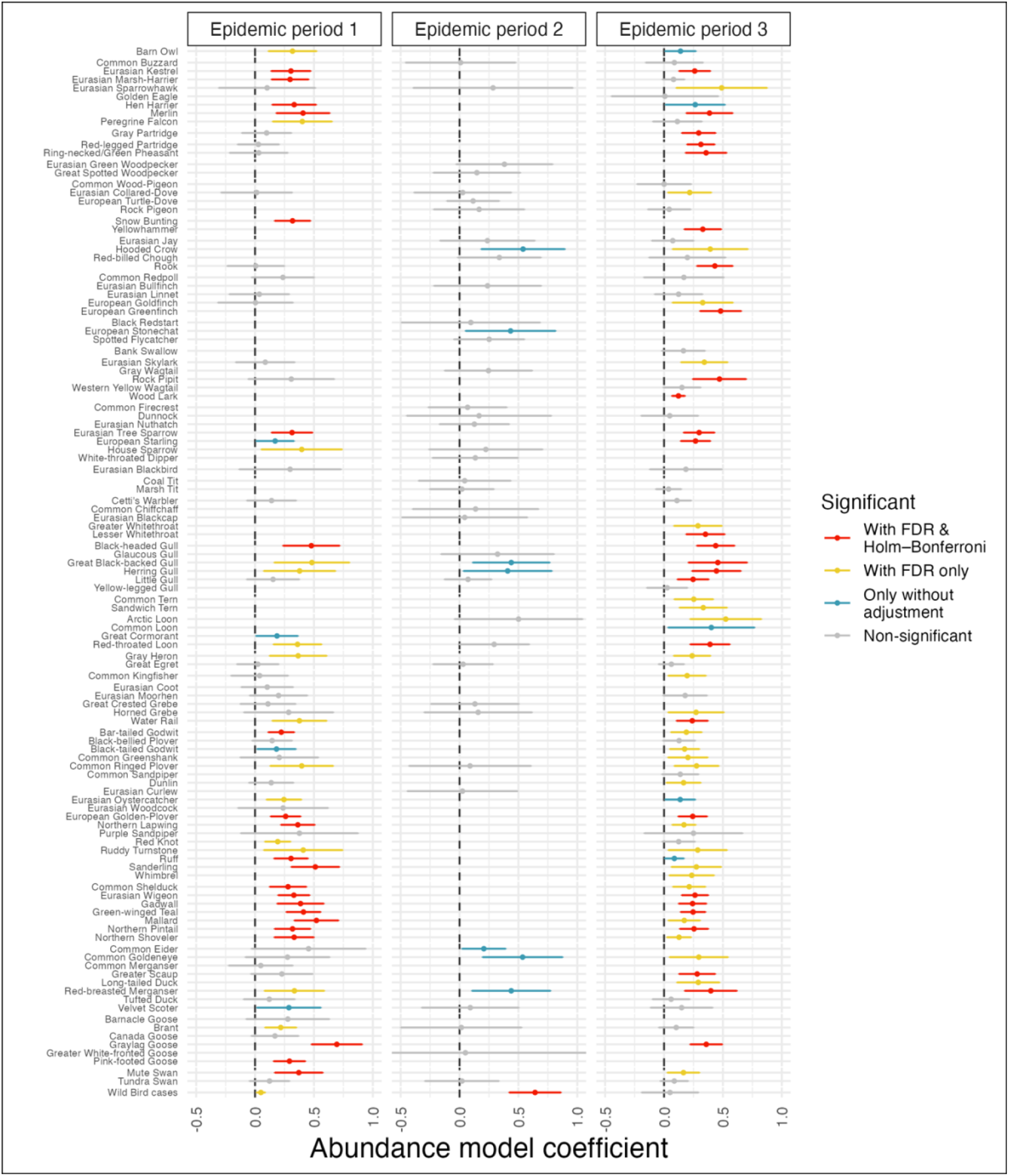
Linear model coefficients for effects of weighted relative species abundance on probability of HPAI cases in premises. ‘Wild Bird cases’ refers to a count of HPAI positive cases in wild birds within the relevant epidemic period rather than weighted relative species abundance. Only positive model coefficients are shown. Error bars indicate the 95% confidence interval. Model coefficients were derived from independent spatial GAM models run for each species and epidemic period (EP1 = 19^th^ October 2021 – 9^th^ February 2022; EP2 = 10^th^ February 2022 – 16^th^ August 2022; EP3 = 17^th^ August 2022 – 20^th^ January 2023).

**S5 Fig.**
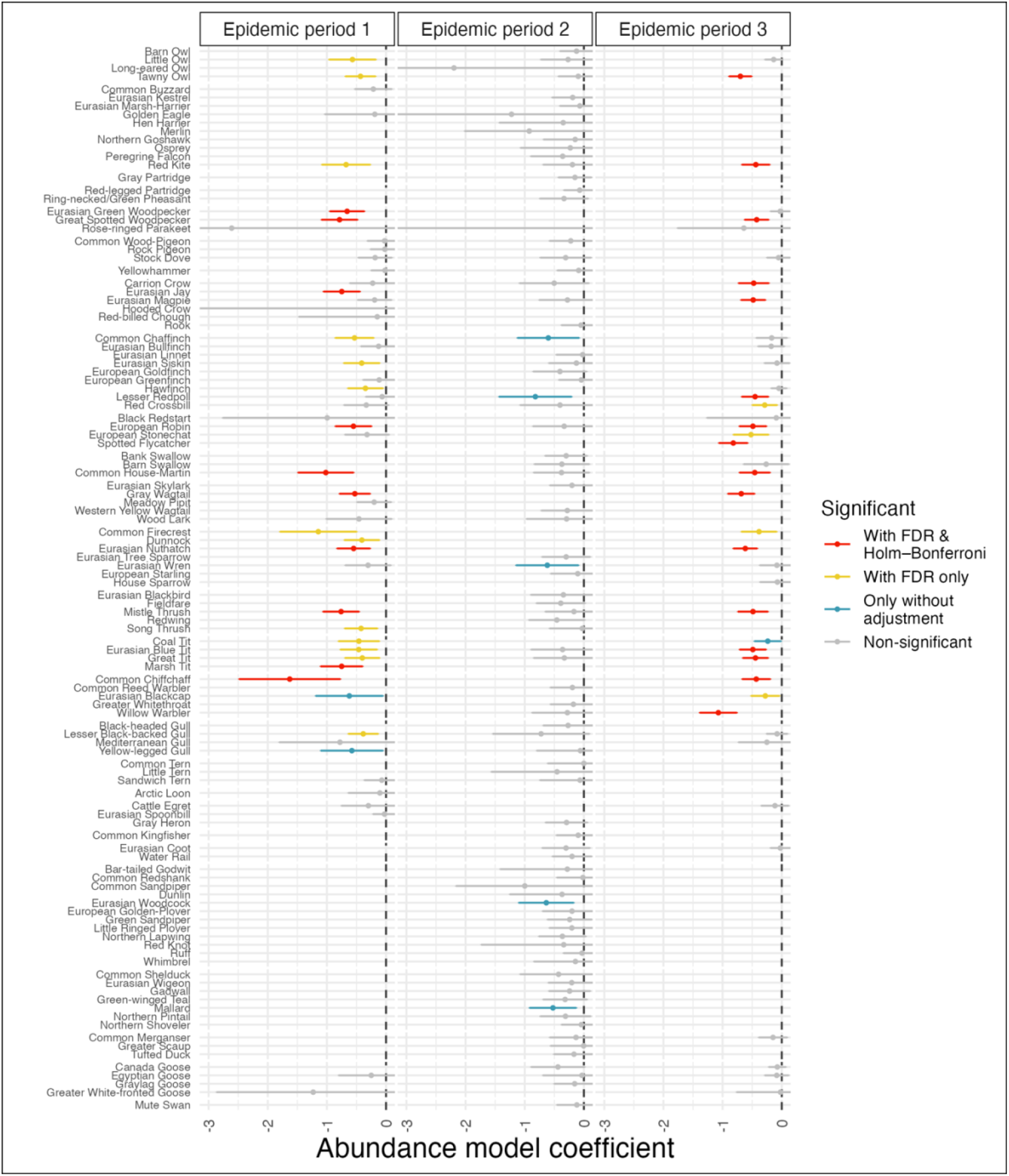
Linear model coefficients for effects of weighted relative species abundance on probability of HPAI cases in premises. ‘Wild Bird cases’ refers to a count of HPAI positive cases in wild birds within the relevant period rather than weighted relative species abundance. Only negative model coefficients are shown. Error bars indicate the 95% confidence interval. Model coefficients were derived from independent spatial GAM models run for each species and epidemic period (EP1 = 19^th^ October 2021 – 9^th^ February 2022; EP2 = 10^th^ February 2022 – 16^th^ August 2022; EP3 = 17^th^ August 2022 – 20^th^ January 2023).

**S6 Fig.**
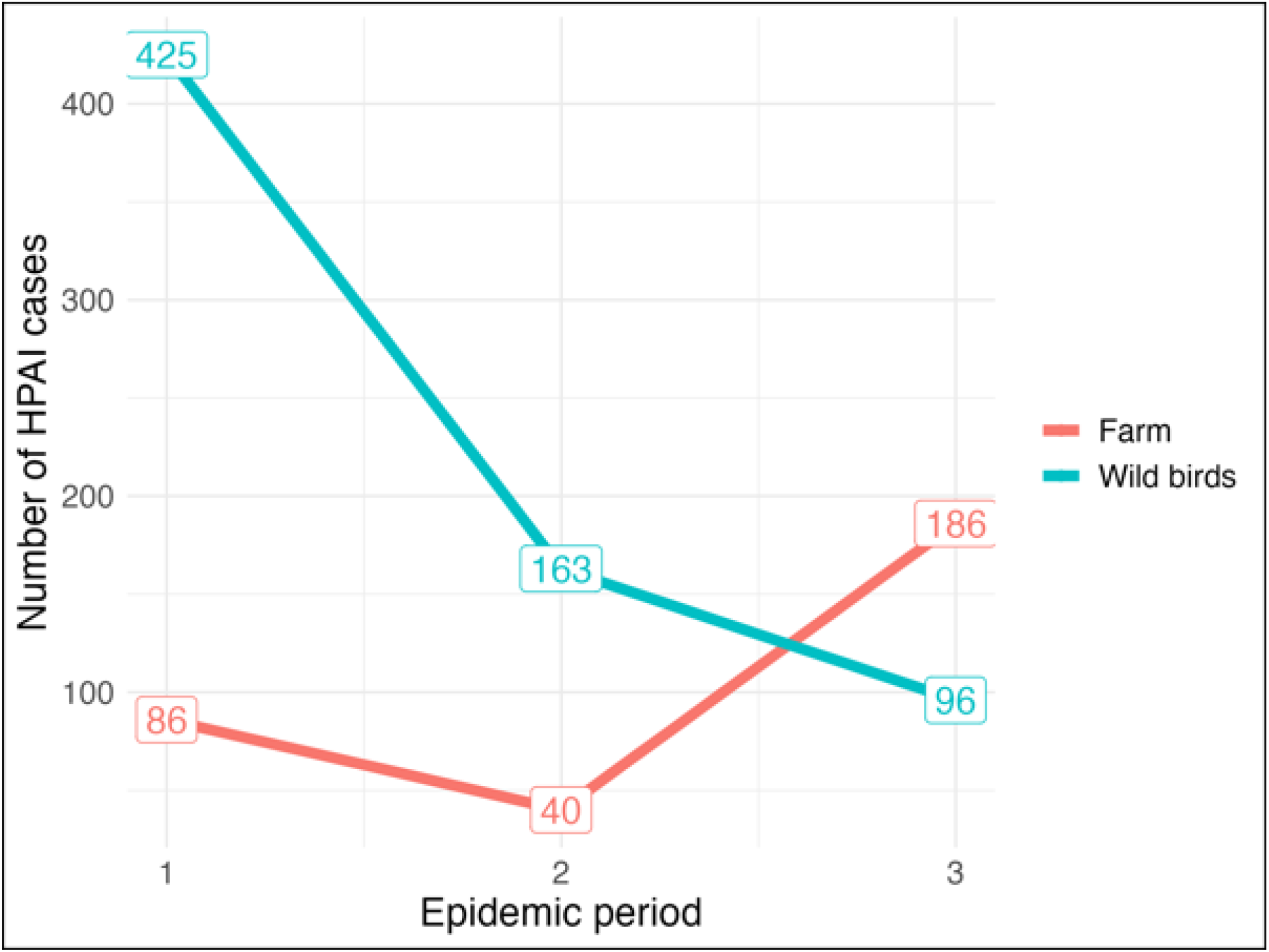
Count of reported cases of highly pathogenic avian influenza (HPAI) across three epidemic periods (EP1 = 19^th^ October 2021 – 9^th^ February 2022; EP2 = 10^th^ February 2022 – 16^th^ August 2022; EP3 = 17^th^ August 2022 – 20^th^ January 2023) on premises (red) and in wild birds (blue). Where multiple individual birds are infected at the same location at the same time this represents a single case.

**S7 Fig.**
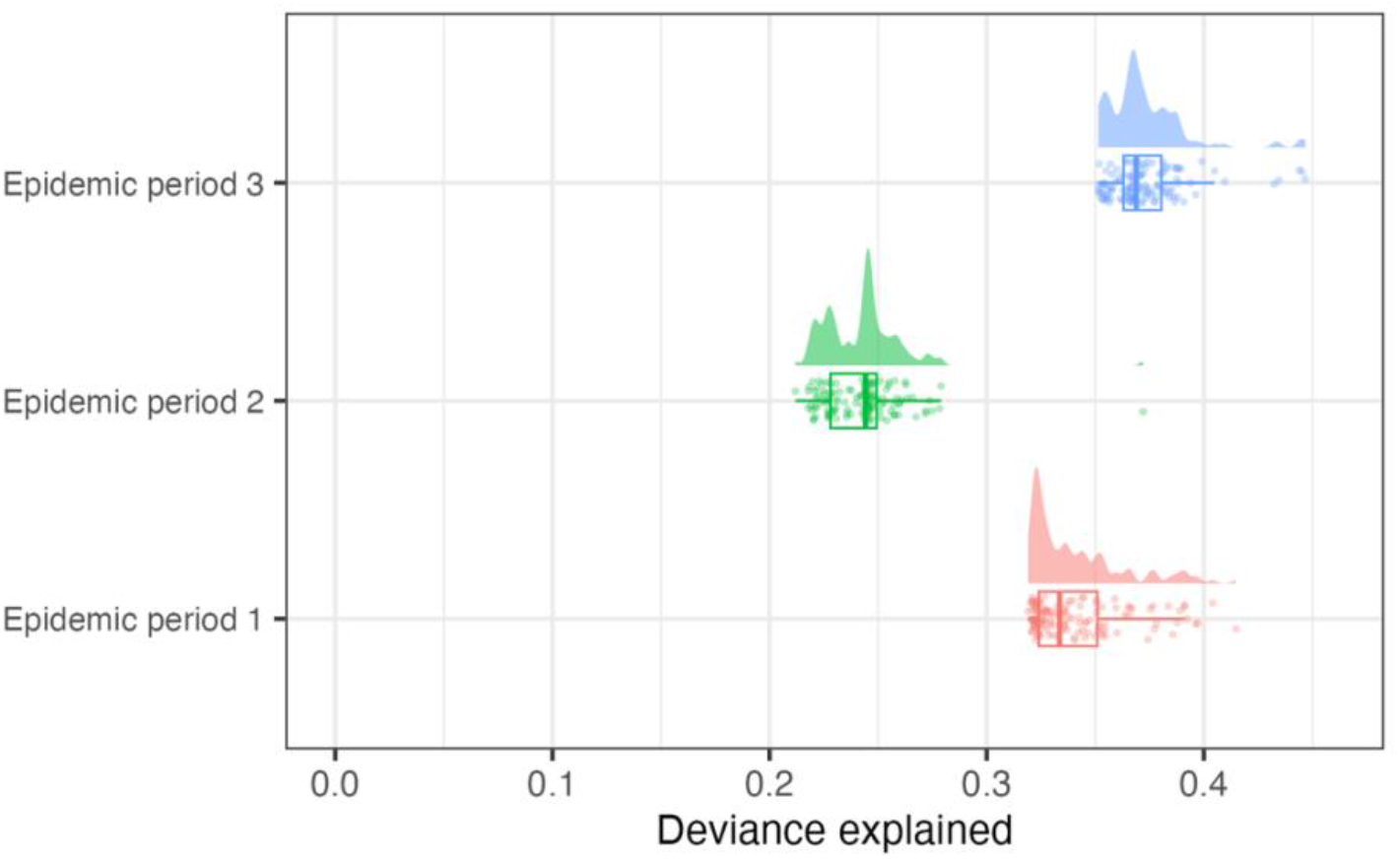
Adjusted R-squared values for each of the 389 independent spatial-GAM models assessing associations between weighted relative species abundance and HPAI cases in premises for each species and epidemic period (EP1 = 19^th^ October 2021 – 9^th^ February 2022; EP2 = 10^th^ February 2022 – 16^th^ August 2022; EP3 = 17^th^ August 2022 – 20^th^ January 2023).

*S1 Data. Linear model coefficients for the intercept, and effects of weighted relative species abundance and total poultry stock numbers on probability of HPAI cases in premises. ‘Wild Bird cases’ refers to a count of HPAI positive cases in wild birds within the relevant period rather than weighted relative species abundance. Model coefficients were derived from independent spatial GAM models run for each species and epidemic period (EP1 = 19^th^ October 2021 – 9^th^ February 2022; EP2 = 10^th^ February 2022 – 16^th^ August 2022; EP3 = 17^th^ August 2022 – 20^th^ January 2023). Species codes are those used by eBird*.

## Literature Cited

Abolnik C, Phiri T, Peyrot B, de Beer R, Snyman A, Roberts D, et al. The Molecular Epidemiology of Clade 2.3.4.4B H5N1 High Pathogenicity Avian Influenza in Southern Africa, 2021–2022. Viruses. 2023;15(6): 1383.

Adlhoch C, Baldinelli F. Avian influenza, new aspects of an old threat. Eurosurveillance. 2023; 28(19): 2300227.

Banyard AC, Lean FZ, Robinson C, Howie F, Tyler G, Nisbet C, et al. Detection of highly pathogenic avian influenza virus H5N1 clade 2.3. 4.4 b in great skuas: a species of conservation concern in Great Britain. Viruses. 2022; 14(2): 212.

Benjamini Y, Hochberg Y. Controlling the false discovery rate: a practical and powerful approach to multiple testing. Journal of the Royal statistical society: series B (Methodological). 1995; 57(1): 289–300.

Blackburn TM, Gaston KJ. Abundance, biomass and energy use of native and alien breeding birds in Britain. Biological invasions. 2018 Dec;20(12):3563–73.

Blagodatski A, Trutneva K, Glazova O, Mityaeva O, Shevkova L, Kegeles E, et al. Avian influenza in wild birds and poultry: dissemination pathways, monitoring methods, and virus ecology. Pathogens. 2021; 10(5): 630.

Bonneaud C, Longdon B. Emerging pathogen evolution: using evolutionary theory to understand the fate of novel infectious pathogens. EMBO reports. 2020; 21(9): e51374.

British Ornithologists Union (BOU). The official list of birds recorded in Britain. 2023. Available at: https://bou.org.uk/british-list/

Byrne AM, James J, Mollett BC, Meyer SM, Lewis T, Czepiel M, et al. Investigating the Genetic Diversity of H5 Avian Influenza Viruses in the United Kingdom from 2020–2022. Microbiology Spectrum. 2023: e04776–22.

Caliendo V, Kleyheeg E, Beerens N, Camphuysen KC, Cazemier R, Elbers AR, Fouchier RA, Kelder L, Kuiken T, Leopold M, Slaterus R. Effect of 2020-21 and 2021-22 Highly Pathogenic Avian Influenza H5 Epidemics on Wild Birds, the Netherlands. Emerging infectious diseases.;30(1).

Clemmons EA, Alfson KJ, Dutton III JW. Transboundary animal diseases, an overview of 17 diseases with potential for global spread and serious consequences. Animals. 2021; 11(7): 2039.

Department for Environment, Food and Rural Affairs (DEFRA). Risk Assessment on the spread of High Pathogenicity Avian Influenza (HPAI) H5N1 to wild birds from released, formerly captive gamebirds in Great Britain: Pheasants. Qualitative Risk Assessment. 2022. Available from: https://assets.publishing.service.gov.uk/government/uploads/system/uploads/attachment_data/file/1124975/Risk_Assessment_on_the_spread_of_High_Pathogenicity_Avian_InfluenzaHPAIH5N1_to_wild_birds_from_releasedformerly_captive_gamebirds_in_Great_Britain_Pheasants.pdf

Department for Environment, Food and Rural Affairs (DEFRA). Updated Outbreak Assessment #39. Highly pathogenic avian influenza (HPAI) in the UK and Europe. 2023A. Available from: https://www.gov.uk/government/collections/animal-diseases-international-monitoring

Department for Environment, Food and Rural Affairs (DEFRA). Highly pathogenic avian influenza H5N1 rapid risk assessment. Catching-up of wild gamebirds in winter 2022 to 2023. 2023B. Available from: https://assets.publishing.service.gov.uk/government/uploads/system/uploads/attachment_data/file/1127836/gamebirds-catching-up-risk-assessment-2022.pdf

Department for Environment, Food and Rural Affairs (DEFRA). High pathogenicity avian influenza (HPAI) in the UK and Europe. Updated Outbreak Assessment #44. 2023C. Available from: https://assets.publishing.service.gov.uk/government/uploads/system/uploads/attachment_data/file/1180257/HPAI_Europe44_18_August_2023_.pdf

Empres-i. Global Animal Disease Information System. 2023. Available from: https://empres-i.apps.fao.org/diseases

European Food Safety Authority (EFSA), Aznar I, Baldinelli F, Stoicescu A, Kohnle L. Annual report on surveillance for avian influenza in poultry and wild birds in Member States of the European Union in 2021. EFSA Journal. 2022 Sep;20(9):e07554.

European Food Safety Authority (EFSA), European Centre for Disease Prevention and Control, European Union Reference Laboratory for Avian Influenza, Adlhoch C, Fusaro A, Gonzales JL, et al. Avian influenza overview April–June 2023. EFSA Journal. 2023A; 21(7): e08191.

European Food Safety Authority (EFSA), European Centre for Disease Prevention and Control, European Union Reference Laboratory for Avian Influenza, Adlhoch C, Fusaro A, Gonzales JL, Kuiken T, Marangon S, Mirinaviciute G, Niqueux É, Stahl K. Avian influenza overview december 2022–march 2023B. EFSA Journal. 2023 Mar;21(3):e07917.

Ewald JA, Sotherton NW, Aebischer NJ. Research into practice: Gray partridge (Perdix perdix) restoration in Southern England. Frontiers in Ecology and Evolution. 2020 Nov 26;8:517500.

Falchieri M, Reid SM, Ross CS, James J, Byrne AM, Zamfir M, et al. Shift in HPAI infection dynamics causes significant losses in seabird populations across Great Britain. Veterinary Record. 2022 Oct;191(7):294–6.

Fink D, Damoulas T, Dave J. Adaptive Spatio-Temporal Exploratory Models: Hemisphere-wide species distributions from massively crowdsourced eBird data. Proceedings of the AAAI Conference on Artificial Intelligence. 2013; 27(1): 1284–1290.

Fink D, Auer T, Johnston A, Ruiz-Gutierrez V, Hochachka WM, Kelling S. Modeling avian full annual cycle distribution and population trends with citizen science data. Ecological Applications. 2020; 30(3): e02056.

Fink D, Auer T, Johnston A, Strimas-Mackey M, Ligocki S, Robinson O, et al. eBird Status and Trends, Data Version: 2022; Released: 2023. Cornell Lab of Ornithology, Ithaca, New York. Available from: 10.2173/ebirdst.2021

Fujiwara M, Auty H, Brown I, Boden L. Assessing the Likelihood of High Pathogenicity Avian Influenza Incursion into the Gamebird Sector in Great Britain via Designated Hatcheries. Frontiers in Veterinary Science. 2022; 9: 877197.

Geen GR, Robinson RA, Baillie SR. Effects of tracking devices on individual birds–a review of the evidence. Journal of Avian Biology. 2019; 50(2).

Gilbert M, Nicolas G, Cinardi G, Van Boeckel TP, Vanwambeke SO, Wint GR, Robinson TP. Global distribution data for cattle, buffaloes, horses, sheep, goats, pigs, chickens, and ducks in 2010. Scientific data. 2018 Oct 30;5(1):1–1.

Gray A, Loeb J. Calls for gamebird release to be paused. The Veterinary Record. 2022; 191(5): 189–189.

Hill NJ, Bishop MA, Trovão NS, Ineson KM, Schaefer AL, Puryear WB, et al. Ecological divergence of wild birds drives avian influenza spillover and global spread. PLoS Pathogens. 2022; 18(5): e1010062.

Hill EM, House T, Dhingra MS, Kalpravidh W, Morzaria S, Osmani MG, et al. The impact of surveillance and control on highly pathogenic avian influenza outbreaks in poultry in Dhaka division, Bangladesh. PLoS computational biology. 2018; 14(9): e1006439.

Hill A, Gillings S, Berriman A, Brouwer A, Breed AC, Snow L, et al. Quantifying the spatial risk of Avian Influenza introduction into British poultry by wild birds. Scientific Reports. 2019; 9(1): 19973.

Holm S. A simple sequentially rejective multiple test procedure. Scandinavian journal of statistics. 1979: 65–70.

Houston DD, Azeem S, Lundy CW, Sato Y, Guo B, Blanchong JA, Gauger PC, Marks DR, Yoon KJ, Adelman JS. Evaluating the role of wild songbirds or rodents in spreading avian influenza virus across an agricultural landscape. PeerJ. 2017 Dec 13;5:e4060.

James J, Seekings AH, Skinner P, Purchase K, Mahmood S, Brown IH, et al. Rapid and sensitive detection of high pathogenicity Eurasian clade 2.3. 4.4 b avian influenza viruses in wild birds and poultry. Journal of Virological Methods. 2022 Mar 1;301:114454.

Kaplan BS, Webby RJ. The avian and mammalian host range of highly pathogenic avian H5N1 influenza. Virus Research. 2013; 178(1): 3–11.

Keawcharoen J, Van Riel D, van Amerongen G, Bestebroer T, Beyer WE, Van Lavieren R, et al. Wild ducks as long-distance vectors of highly pathogenic avian influenza virus (H5N1). Emerging infectious diseases. 2008 Apr;14(4):600.

Kim JK, Negovetich NJ, Forrest HL, Webster RG. Ducks: the “Trojan horses” of H5N1 influenza. Influenza and other respiratory viruses. 2009; 3(4): 121–8.

Kou Z, Lei FM, Yu J, Fan ZJ, Yin ZH, Jia CX, et al. New genotype of avian influenza H5N1 viruses isolated from tree sparrows in China. Journal of virology. 2005; 79(24): 15460–6.

Lane JV, Jeglinski JW, Avery-Gomm S, Ballstaedt E, Banyard AC, Barychka T, et al. High pathogenicity avian influenza (H5N1) in Northern Gannets: Global spread, clinical signs, and demographic consequences. Ibis. 2023.

Lee DH, Bertran K, Kwon JH, Swayne DE. Evolution, global spread, and pathogenicity of highly pathogenic avian influenza H5Nx clade 2.3. 4.4. Journal of veterinary science. 2017; 18(S1): 269–80.

Lewis NS, Banyard AC, Whittard E, Karibayev T, Al Kafagi T, Chvala I, et al. Emergence and spread of novel H5N8, H5N5 and H5N1 clade 2.3. 4.4 highly pathogenic avian influenza in 2020. Emerging Microbes & Infections. 2021; 10(1): 148–51.

Liang Y, Hjulsager CK, Seekings AH, Warren CJ, Lean FZ, Núñez A, Jet al. Pathogenesis and infection dynamics of high pathogenicity avian influenza virus (HPAIV) H5N6 (clade 2.3. 4.4 b) in pheasants and onward transmission to chickens. Virology. 2022; 577: 138–48.

Lupiani B, Reddy SM. The history of avian influenza. Comparative Immunology, Microbiology and Infectious Diseases. 2009; 32(4): 311–23.

Madden JR. How many gamebirds are released in the UK each year?. European Journal of Wildlife Research. 2021; 67(4): 72.

Madden JR, Hall A, Whiteside MA. Why do many pheasants released in the UK die, and how can we best reduce their natural mortality?. European Journal of Wildlife Research. 2018 Aug;64:1–3.

Martin V, Dobschuetz SV, Lemenach A, Rass N, Schoustra W, DeSimone L. Early warning, database, and information systems for avian influenza surveillance.

Pearce-Higgins JW, Humphreys EM, Burton NH, Atkinson PW, Pollock C, Clewley GD, Johnston DT, O’Hanlon NJ, Balmer DE, Frost TM, Harris SJ. Highly pathogenic avian influenza in wild birds in the United Kingdom in 2022: impacts, planning for future outbreaks, and conservation and research priorities. Report on virtual workshops held in November 2022. BTO Research Report. 2023;752.

Pohlmann A, Stejskal O, King J, Bouwhuis S, Packmor F, Ballstaedt E, et al. Mass mortality among colony-breeding seabirds in the German Wadden Sea in 2022 due to distinct genotypes of HPAIV H5N1 clade 2.3.4.4b. Journal of General Virology. 2023; 104(4): 001834.

Portier J, Ryser-Degiorgis MP, Hutchings MR, Monchâtre-Leroy E, Richomme C, Larrat S, et al. Multi-host disease management: the why and the how to include wildlife. BMC veterinary research. 2019; 15(1): 1–1.

Poulson RL, Brown JD. Wild bird surveillance for avian influenza virus. Animal Influenza Virus: Methods and Protocols. 2020: 93–112.

Ramey AM, Hill NJ, DeLiberto TJ, Gibbs SE, Camille Hopkins M, Lang AS, et al. Highly pathogenic avian influenza is an emerging disease threat to wild birds in North America. The Journal of Wildlife Management. 2022; 86(2): e22171.

R Core Team. R: A language and environment for statistical computing. R Foundation for Statistical Computing, Vienna, Austria. 2023. Available at: https://www.R-project.org/.

Sánchez-Cano A, Camacho MC, Ramiro Y, Cardona-Cabrera T, Höfle U. Seasonal changes in bird communities on poultry farms and house sparrow—wild bird contacts revealed by camera trapping. Frontiers in Veterinary Science. 2024 Feb 20;11:1369779.

Seekings AH, Warren CJ, Thomas SS, Lean FZ, Selden D, Mollett BC, et al. Different Outcomes of Chicken Infection with UK-Origin H5N1-2020 and H5N8-2020 High-Pathogenicity Avian Influenza Viruses (Clade 2.3. 4.4 b). Viruses. 2023 Sep 12;15(9):1909.

Slomka MJ, Reid SM, Byrne AM, Coward VJ, Seekings J, Cooper JL, et al. Efficient and Informative Laboratory Testing for Rapid Confirmation of H5N1 (Clade 2.3. 4.4) High-Pathogenicity Avian Influenza Outbreaks in the United Kingdom. Viruses. 2023; 15(6): 1344.

Strimas-Mackey M, Ligocki S, Auer T, & Fink D. ebirdst: Tools for loading, plotting, mapping and analysis of eBird Status and Trends data products. R package version 2.2021.1. 2022. Available from: https://ebird.github.io/ebirdst/.

Uher-Koch BD, Spivey TJ, Van Hemert CR, Schmutz JA, Jiang K, Wan XF, et al. Serologic Evidence for Influenza a Virus Exposure in Three Loon Species Breeding in Alaska, USA. Journal of wildlife diseases. 2019 Oct 1;55(4):862–7.

Viana M, Mancy R, Biek R, Cleaveland S, Cross PC, Lloyd-Smith JO, et al. Assembling evidence for identifying reservoirs of infection. Trends in ecology & evolution. 2014; 29(5): 270–9.

Wade D, Ashton-Butt A, Scott G, Reid SM, Coward V, Hansen RD, Banyard AC, Ward AI. High pathogenicity avian influenza: targeted active surveillance of wild birds to enable early detection of emerging disease threats. Epidemiology & Infection. 2023;151:e15.

Wille M, Lisovski S, Roshier D, Ferenczi M, Hoye BJ, Leen T, et al. Strong host phylogenetic and ecological effects on host competency for avian influenza in Australian wild birds. Proceedings of the Royal Society B. 2023; 290(1991): 20222237.

Wood SN. Fast stable restricted maximum likelihood and marginal likelihood estimation of semiparametric generalized linear models. Journal of the Royal Statistical Society Series B: Statistical Methodology. 2011; 73(1): 3–6.

